# Capsaicin acts as a novel NRF2 agonist to suppress ethanol induced gastric mucosa oxidative damage by directly disrupting the KEAP1-NRF2 interaction

**DOI:** 10.1101/2024.01.06.574490

**Authors:** Xiaoning Gao, WuYan Guo, Peiyuan Liu, Mingyue Yuwen, Hongyu Ren, Shengtao Hu, Zixiang Liu, Ruyang Tan, Kairui Liu, Zhiru Yang, Junli Ba, Xue Bai, Shiti Shama, Cong Tang, Kai Miao, Haozhi Pei, Liren Liu, Cheng Zhu, Tao Wang, Bo Zhang, Jun Kang

## Abstract

Excessive alcohol consumption poses significant health risks and is closely associated with oxidative damage. The KEAP1-NRF2-ARE signaling pathway serves as the primary antioxidant system. However, current small molecule inhibitors are all covalently bound to KEAP1, meaning that once bound, they are not easily dissociated, while continuous inhibition of KEAP1 exhibits severe side effects. In this study, BLI, CETSA, Pull-down, Co-IP and HDX-MS assay analysis were conducted to detect the KEAP1 binding behavior of natural product, capsaicin (CAP), both *in vitro* and in cells. The ethanol-induced acute gastric mucosal damage rat model was also established to evaluate the therapeutic effect of CAP. Our findings demonstrated that CAP mitigated mitochondrial damage, facilitated the nuclear translocation of NRF2, leading to the up-regulation of downstream antioxidant response elements, HMOX1, TXN, GSS and NQO1 in GES-1 cells. Furthermore, CAP directly bind to KEAP1 and inhibit the interaction between KEAP1 and NRF2. In the KEAP1-knockout 293T cells, CAP failed to activate NRF2 expression. We identified that CAP non-covalently bound to Kelch domain and allosterically regulated three specific regions of KEAP1 : L342-L355, D394-G423 and N482-N495. To improve drug solubility and delivery efficiency, we developed IR-Dye800 modified albumin coated CAP nanoparticles. The nanoparticles significantly reduced the gastric mucosal inflammation and activated NRF2 downstream genes *in vivo*. Our hypothesis was further verified our hypothesis in Nfe2l2-knockout mice. This study provides new insights that CAP is a safe and novel NRF2 agonist by allosterically regulating KEAP1, which may contribute to the development of lead drugs for oxidative stress-related illness, e.g. aging, cancer, neurodegenerative and cardiovascular diseases.

## 1 Introduction

Ethanol (EtOH), commonly referred to alcohol in everyday life, is a fascinating ingredient found in beer, wine, and liquor. Excessive consumption poses great health risks and is responsible for approximately 3 million deaths per year^1^. In 2016, the largest share of all deaths caused by alcohol consumption worldwide was attributed to digestive disorders (21.3%)^1^. Alcohol induces oxidative damage to gastric mucosa. Persistent and intensified oxidative stress, particularly reactive oxygen species (ROS), induces DNA damage, protein oxidation, and lipid peroxidation^2^. Consequently, this can result in gastrointestinal dysfunction, chronic atrophic gastritis, and is closely related to the development of gastric cancer^3^.

Natural products are essential sources of drugs or lead compounds for treating major diseases^4^.Previous reports indicate that the incidence rate of gastric ulcers in Indians and Malaysians is quite low. Chili peppers are commonly found in the cuisine of these two countries^5^. This raises questions regarding the potential for chili peppers to alleviate gastric mucosal damage and reduce the occurrence of gastric ulcers. Recent experiments conducted in rats have demonstrated that red pepper/capsaicin (CAP) possesses significant protective effects on ethanol-induced gastric mucosal damage, and the mechanisms involved may relate to the promotion of vasodilation^6,7^, increased mucus secretion^8^ and the release of calcitonin gene-related peptide (CGRP)^9, 10^. However, it is important to note that the specific role of the antioxidant activity of CAP has not been thoroughly investigated.

The Kelch-like ECH-associated protein 1 (KEAP1)–Nuclear factor erythroid 2–related factor 2 (NRF2)–antioxidant response element (ARE) pathway is a core defense mechanism against oxidative and electrophilic stress^11^. Under homeostatic conditions, KEAP1 acts as a linker protein for the Cul3-E3 ubiquitin ligase complex, continuously promoting the ubiquitination and proteasomal degradation of NRF2, thereby maintaining NRF2 at basal levels^12^. When oxidative or electrophilic stress occurs, critical cysteine residues in KEAP1 are modified, or the interaction between the ETGE/DLG motifs on NRF2 and the Kelch domain of KEAP1 is disrupted, allowing NRF2 to escape degradation, accumulate, and translocate to the nucleus. There, NRF2 forms heterodimers with small Maf proteins and binds to ARE, inducing the expression of antioxidant and cytoprotective genes such as those involved in glutathione metabolism, NADPH regeneration, phase II detoxifying enzymes, and drug efflux transporters, thereby restoring redox balance within the cell and reducing oxidative damage^13^.

Classical NRF2 agonists, such as sulforaphane, are small molecules that bind to KEAP1 and covalently modify its cysteine residues, thereby altering the binding affinity between KEAP1 and NRF2 ^14^. However, traditional covalent agonists may induce sustained overactivation of NRF2, leading to adverse side effects and limiting clinical application ^15^. Consequently, recent efforts have shifted toward the development of non-covalent NRF2 agonists, which are generally associated with lower toxicity and greater translational potential, enabling more controlled enhancement of NRF2 activity and offering new insights and therapeutic opportunities in antioxidant-related interventions.

In this study, EtOH induced acute oxidative damage models were established *in vitro* and *in vivo*, and the effects of CAP pretreatment on KEAP1-NRF2-ARE signaling pathway were verified. Ultimately, we employed hydrogen-deuterium exchange mass spectrometry (HDX-MS) to identify the allosteric regulatory sites following CAP treatment. CAP may emerge as a pioneering direct inhibitor of the KEAP1-NRF2 interaction, which also offers an in-depth understanding of its antioxidative defense mechanism.

## 2 Results

### 2.1 Capsaicin protects GES-1 from oxidative stress

The molecular formula of CAP (PubChem CID: 1548943) is provided in **Supplemental Figure 1a**. Initially, an oxidative stress model was established in human gastric mucosal epithelial (GES-1) cells. According to the *Lancet*^16^, WHO recommendation and US Standard Drink Sizes, that daily alcohol consumption should be limited to 1-2 standard cups/day (1 standard cup is equivalent to 10-14 g of pure EtOH in different countries), so cell models were tested at a similar concentration of 5%. A dose-dependent decline in cell viability upon exposure to EtOH was observed, with viability decreasing to approximately 50% after 1.5 hours of treatment with 5% EtOH **(Supplemental Figure 1b)**. To evaluate the protective effects of CAP, GES-1 model cells were pretreated with CAP for 2.5 hours before introducing EtOH. The results revealed that a pretreatment with 8 μM CAP significantly enhanced cell viability to 93.84% **(Supplemental Figure 1c)**. Notably, further increases in CAP concentration did not enhance the protective effect, possibly due to cytotoxicity at higher concentrations^17^**(Supplemental Figure 1d)**. Subsequently, three CAP concentrations (2, 8, and 32 μM) and three EtOH concentrations (0.5%, 3.5%, and 5%) on GES-1 were further valuated to establish a stable cell model *in vitro* **(Supplemental Figure 1e-g)**. Furthermore, we assessed EtOH-induced apoptosis and found that the percentage of apoptotic cells was reduced from 28.85% to 13.95% in the group pretreated with 8 μM CAP, compared to the EtOH-only group **(Supplemental Figure 1h)**.

Then the morphological changes in GES-1 cells were examined. Cells in the EtOH treated group displayed a rounded morphology and showed a marked tendency for aggregation. In contrast, cells that were pretreated with 2 μM and 8 μM CAP largely retained their characteristic cellular shape **(Figure 1a)**. Then, we assessed ROS production using DCFH-DA staining. The EtOH-only group exhibited elevated levels of ROS, as indicated by increased green fluorescence, while the phenomenon was substantially diminished in CAP-pretreated groups **(Figure 1b)**. Flow cytometric analyses revealed a consistent decline in intracellular ROS levels with increasing dosages of CAP: from 83.1±3.59% in the ethanol-only group to 60.23±12.67% and 48.9±5.55% in the 2 μM and 8 μM CAP groups, respectively **(Figure 1c and 1d)**. To evaluate the impact of CAP on intracellular redox homeostasis, the activity of superoxide dismutase (SOD), as well as the amount of malondialdehyde (MDA) were measured. Our findings demonstrated that SOD levels were significantly upregulated following CAP treatment **(Figure 1e)**, while MDA production was markedly reduced **(Figure 1f)**. Collectively, these results suggest that CAP effectively reduces ROS levels and enhances redox balance in GES-1 cells exposed to EtOH.

**Figure 1.**
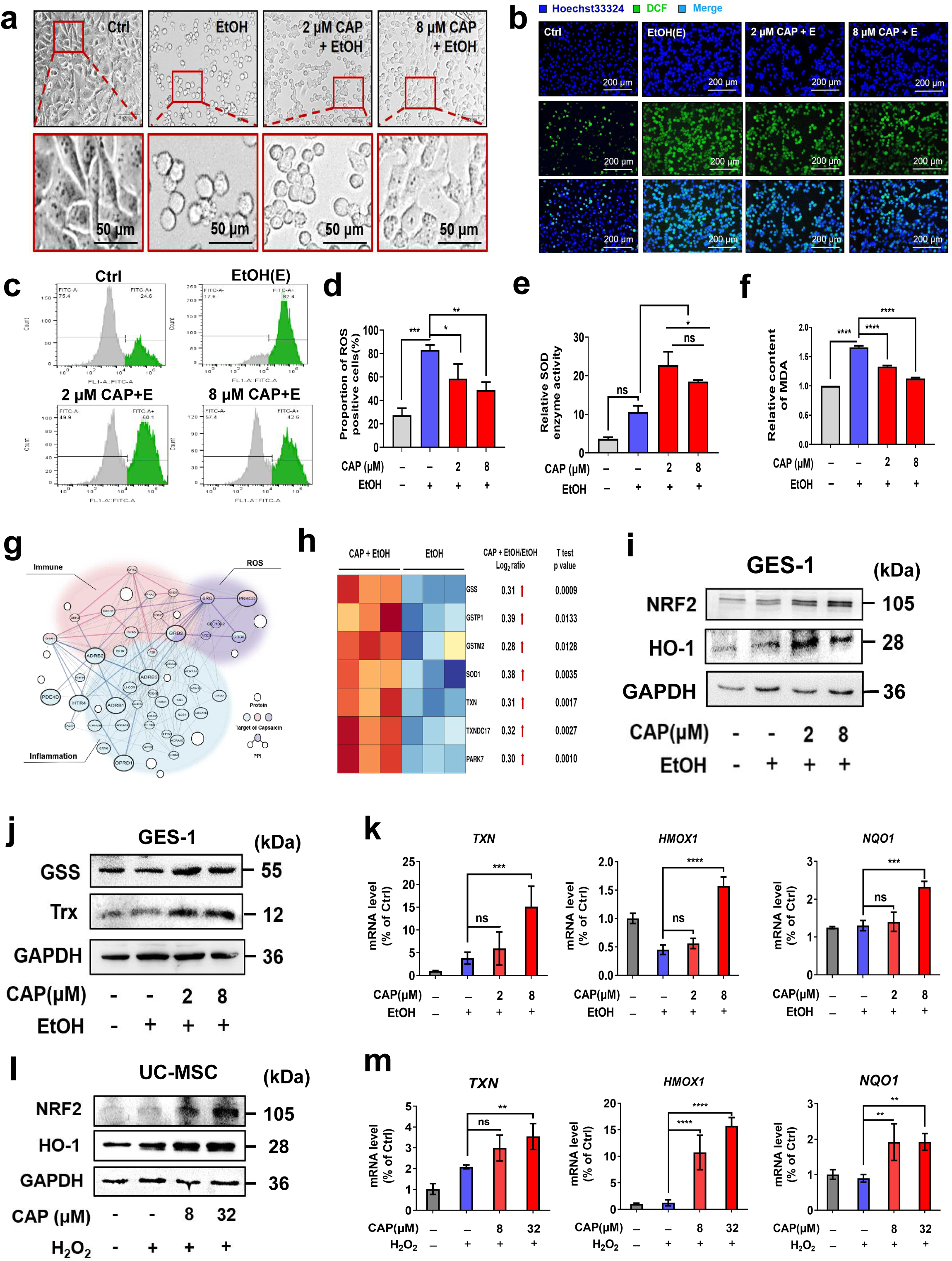
Capsaicin reduces ROS and promotes NRF2 expression. **(a)** Microscopic examination of GES-1 cellular morphology following treatment regimens. Cells were pre-treated with Capsaicin (CAP) at concentrations of 2 or 8 μM for 2.5 hours, followed by incubation with 5% EtOH for 1.5 hours. Scale bar, 50 μm. **(b)** Detection of ROS-positive GES-1 cells using DCFH-DA staining. Scale bar, 200 μm. **(c)** Flow cytometric (FCM) analysis of ROS-positive GES-1 cells labeled with DCFH-DA. **(d)** Statistical analysis of ROS levels measured by FCM. The experiment was conducted in triplicate. **(e)** Assessment of SOD activity in GES-1 cells. **(f)** Quantification of MDA levels of GES-1 cells. **(g)** A network targets of CAP. Purple, blue, pink nodes represent proteins or genes in the predicted biological effect profile of CAP related to ROS, inflammation or immune, respectively. **(h)** Heatmap depicting differentially expressed genes (DEGs) enriched in antioxidant activity based on proteomic analysis. Each row represents the expression level of a single gene, while each column corresponds to an individual experimental sample. **(i)** Western blot analysis of NRF2 and HO-1 expression in GES-1 cells. **(j)** Western blot assessment of antioxidant protein expression in GES-1 cells. **(k)** Quantitative analysis of relative mRNA levels of antioxidant response elements (ARE)-related genes in GES-1 cells. Cells were treated with or without CAP and EtOH, and the mRNA levels of TXN, HMOX1, and NQO1 were measured. Each group consisted of three replicates. **(l)** Western blot analysis of NRF2 and HO-1 expression in human umbilical cord mesenchymal stem cells (HUC-MSCs). **(m)** Quantitative analysis of relative mRNA levels of ARE-related genes in HUC-MSCs. The mRNA levels of TXN, HMOX1, and NQO1 were measured in different treatment groups.

### 2.2 Capsaicin activates the NRF2-ARE signaling pathway

With analytical techniques such as network pharmacology^18^ and single-cell sequencing, researchers identified pivotal target proteins implicated in the precursor lesions of gastric cancer^19^. Based on the network targets analysis, CAP may regulate pathways and biological processes related to ROS, inflammation and immune expression **(Figure 1g)**. However, the specific mechanism of ROS regulation by CAP remains unclear, so we conducted proteomic analyses to identify differentially expressed genes (DEGs). A comprehensive overview of these DEGs was presented in the heat map. Using stringent criteria for statistical significance (|Fold Change| > 1.2, P value < 0.05), analysis revealed that 120 proteins were significantly up-regulated, while 65 were down-regulated following CAP treatment **(Supplementary Figure 2a)**. Subsequent Gene Ontology (GO), and Kyoto Encyclopedia of Genes and Genomes (KEGG) enrichment analyses were performed on this pool of 185 DEGs **(Supplementary Figure 2b and 2c)**. As anticipated, the top terms in the ‘Molecular Function’ category of the GO analysis included ‘antioxidant activity’ (GO: 0016209) and ‘oxidoreductase activity’ (GO: 0016491). Genes corresponding to these functional categories were highlighted in the volcano plot **(Supplemental Figure 2d)** and annotated in the heat map **(Figure 1h)**. Intriguingly, our findings demonstrated a significant elevation in the expression of key antioxidant-related proteins, such as glutathione synthase (GSS), superoxide dismutase 1 (SOD1), and thioredoxin (TXN, also known as Trx). These proteins were associated with previously reported antioxidant response elements (AREs) and function as downstream targets of the transcription factor NRF2^20^. We hypothesize that CAP may activate the NRF2-ARE signaling pathway in oxidative stress models.

To substantiate the hypothesis, we discovered that CAP significantly increased the expression levels of NRF2 and heme oxygenase 1 (HMOX1, also known as HO-1) in GES-1 cells **(Figure 1i)**, which is a downstream target of NRF2. Concurrently, the elevated expression levels of GSS and Trx proteins, which were also downstream targets of NRF2, were further validated by western blotting **(Figure 1j)**. A grayscale quantification of these pivotal proteins was presented in **Supplementary Figure 2e-2h**. Recognizing NRF2’s role as a transcriptional regulator, we further analyzed the transcript levels of these essential target genes: TXN, HMOX1 and NQO1. It’s worth mentioning that NAD(P)H quinone dehydrogenase 1 (NQO1) is also a classic target gene of NRF2^21^. Quantitative reverse transcription PCR (RT-qPCR) assays demonstrated that CAP significantly increased the transcription of these antioxidant-associated genes **(Figure 1k)**.

In a parallel experiment involving a hydrogen peroxide (H_2_O_2_)-induced oxidative stress model in human umbilical cord mesenchymal stem cells (UC-MSC), CAP pretreatment similarly boosted the transcription and protein expression levels of NRF2 and its downstream genes of TXN, HMOX1, and NQO1 **(Figure 1l and 1m)**. A grayscale quantification of these pivotal proteins was also shown in **Supplementary Figure 2i and 2j.** These collective findings suggested that CAP’s role in ROS mitigation was not confined to a specific cell type, needing further exploration of the underlying mechanisms

### 2.3 Capsaicin promotes NRF2 to translocate into nucleus

Furthermore, we proceeded to investigate the subcellular localization of NRF2 following pretreatment with CAP. Our findings revealed that CAP effectively facilitated the translocation of NRF2 into the nucleus. Specifically, 52.29% of NRF2 fluorescence co-localized with the nuclear dye DAPI, as determined by analyses across three randomly selected fields of view for each sample group **(Figure 2a and 2b)**. To corroborate this observation, we conducted a nucleoplasm separation assay to quantify the expression levels of NRF2 in both the nuclear and cytoplasmic compartments **(Figure 2c)**. Grayscale analysis of the immunoblotting data further corroborated that the nuclear accumulation of NRF2 increased with CAP treatment **(Figure 2d)**.

**Figure 2.**
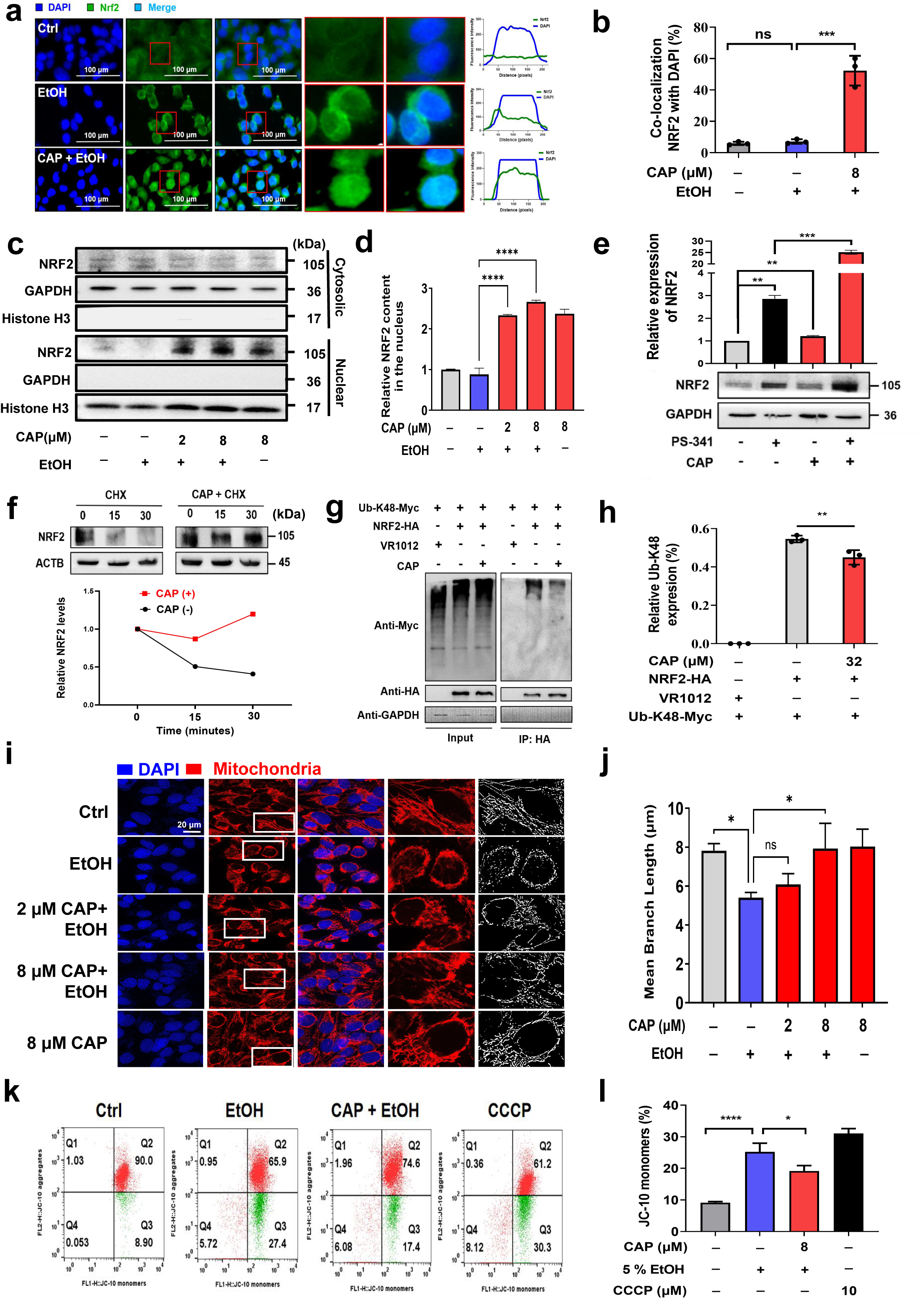
Capsaicin inhibits the ubiquitination and degradation of NRF2. **(a)** Immunofluorescence detection of NRF2 nuclear localization DAPI was employed to label the cell nuclei for reference. Scale bar, 100 μm. **(b)** Statistical analysis of NRF2 nuclear translocation following 8 μM CAP pre-treatment. The proportion of NRF2 localized in the nucleus post-CAP treatment was quantitatively assessed. **(c and d)** Subcellular localization of NRF2 in GES-1 cells across different treatment groups. NRF2 levels in both the nucleus and cytoplasm were assessed. GAPDH was used as a cytoplasmic marker, and Histone H3 served as a nuclear marker. Statistical analysis was performed specifically on the nuclear localization of NRF2. **(e)** Total NRF2 levels induced by PS-341 or CAP. **(f)** Analysis of total NRF2 levels under the influence of cycloheximide (CHX), With or Without 8 μM CAP treatment. NRF2 degradation was semi-quantitatively assessed using ImageJ software to analyze the western blot results. **(g-h)** Inhibition of K48 ubiquitination on NRF2 protein by 32 μM CAP as assessed by Co-IP assay in 293T cells. **(i)** Mitochondrial visualization in GES-1 cells following CAP and EtOH treatment regimens. Cells were pre-treated with CAP at concentrations of 2 or 8 μM for 2.5 hours, followed by a 10-minute incubation with 5% EtOH. Mitochondria were labeled with Mito Tracker Red CMXRos and detected via the Leica STELLARIS 5 Confocal Microscope Platform. Red fluorescence indicates the mitochondria, while blue fluorescence (DAPI staining) marks the cell nucleus. Scale bar represents 20 μm. **(j)** Quantitative analysis of mitochondrial branch length in different treatment groups using ImageJ and GraphPad Prism. The branch length of individual mitochondria was analyzed using ImageJ software and the data were plotted using GraphPad Prism. **(k)** Assessment of mitochondrial membrane potential (ΔΨm) in GES-1 cells using FCM. Cells were pre-treated with CAP at concentrations of 2 or 8 μM for 2.5 hours, followed by a 10-minute incubation with 5% EtOH or a 20-minute incubation with CCCP as a positive control. ΔΨm was evaluated using JC-10 staining. **(l)** Quantitative analysis of ΔΨm across different treatment groups using ImageJ Software.

### 2.4 Capsaicin inhibits the degradation of NRF2

NRF2 was known to undergo proteasomal degradation, so we employed PS-341, a renowned proteasome inhibitor, to selectively impede the 20S proteasome activity. Our data revealed that PS-341 independently elevated NRF2 expression levels. Moreover, when cells were co-incubated with both PS-341 and CAP, the overall NRF2 levels rose even further, which may suggest that the mechanism of CAP differs from that of PS-341 **(Figure 2e)**.

To delve deeper into whether CAP affects protein degradation, we utilized cycloheximide (CHX), a recognized inhibitor of protein synthesis^22^. As shown in **Figure 2f**, CHX treatment led to a rapid decline in NRF2 protein levels, rendering it nearly undetectable approximately 30 minutes after treatment. Conversely, NRF2 levels remained noticeably stable when co-incubated with CAP, suggesting that CAP prolongs NRF2 half-life. These findings support the idea that CAP enhances NRF2 stability by inhibiting its proteolytic degradation. In alignment with existing literature, which posits that K48 ubiquitination facilitates NRF2 degradation^23, 24^, we carried out ubiquitin immunoprecipitation experiments. These indicated a modest reduction, approximately 17.57%, in K48 ubiquitination of NRF2 following CAP treatment **(Figure 2g and 2h)**. This phenomenon indicated that there may be other ways to inhibit degradation

### 2.5 Capsaicin protects mitochondrial function

The mitochondria serve as the primary intracellular source of ROS^25^. Initial exploratory studies focused on the role of NRF2 in maintaining mitochondrial integrity^26^. Our findings revealed a 21.76% decrease in average mitochondrial branch length after treatment with EtOH. Remarkably, pretreatment with CAP was found to preserve mitochondrial structural integrity in GES-1 cells **(Figure 2i and 2j)**. We subsequently turned our attention to the mitochondrial membrane potential (MMP, ΔΨm), a key factor in sustaining mitochondrial oxidative phosphorylation and the subsequent production of adenosine triphosphate (ATP). Exposure to 5% EtOH for 10 minutes or the positive control CCCP for 20 minutes resulted in comparable declines in ΔΨm, affecting approximately 30% of the cells. Notably, pretreatment with 8 μM CAP significantly mitigated this reduction in ΔΨm **(Figure 2k and 2l)**.

Typically, diminished levels of ATP serve as a hallmark of compromised mitochondrial function. To investigate this, we employed a luminescence detector to quantify both extracellular and intracellular ATP concentrations. The extracellular ATP levels surged to approximately three times in the EtOH-only group compared to the control, while a corresponding sharp decrease was observed in intracellular ATP concentrations **(Supplementary Figure 3a)**. In striking contrast, CAP pretreatment effectively stabilized intracellular ATP levels, maintaining them at 96.65% of their original concentration **(Supplementary Figure 3b)**. Building on these observations, we assessed the end product of the anaerobic glycolytic pathway, lactate (LA), as well as the transcription levels of key metabolic enzymes—pyruvate kinase M2 (PKM2) and lactate dehydrogenase A (LDHA). CAP treatment resulted in significant downregulation of both LA and the aforementioned enzymes, suggesting a potential metabolic reprogramming from glycolysis to oxidative phosphorylation **(Supplementary Figure 3c-3f)**. Previous studies have demonstrated that CAP can directly bind to and inhibit the activity of PKM2 and LDHA, subsequently attenuating inflammatory responses^27^. Therefore, our findings underscore the role of CAP in maintaining mitochondrial function and also reducing oxidative stress.

### 2.6 CAP disrupts KEAP1-NRF2 interaction

To explore the possible influence of CAP on NRF2 stabilization via KEAP1 modulation, our findings revealed no significant alterations in KEAP1 expression, both at the transcriptional level **(Figure 3a)** and at the protein level **(Figure 3b)**. The possibility that CAP exerted no impact on KEAP1 expression suggests the likelihood that CAP may disrupt the interaction between KEAP1 and NRF2.

**Figure 3.**
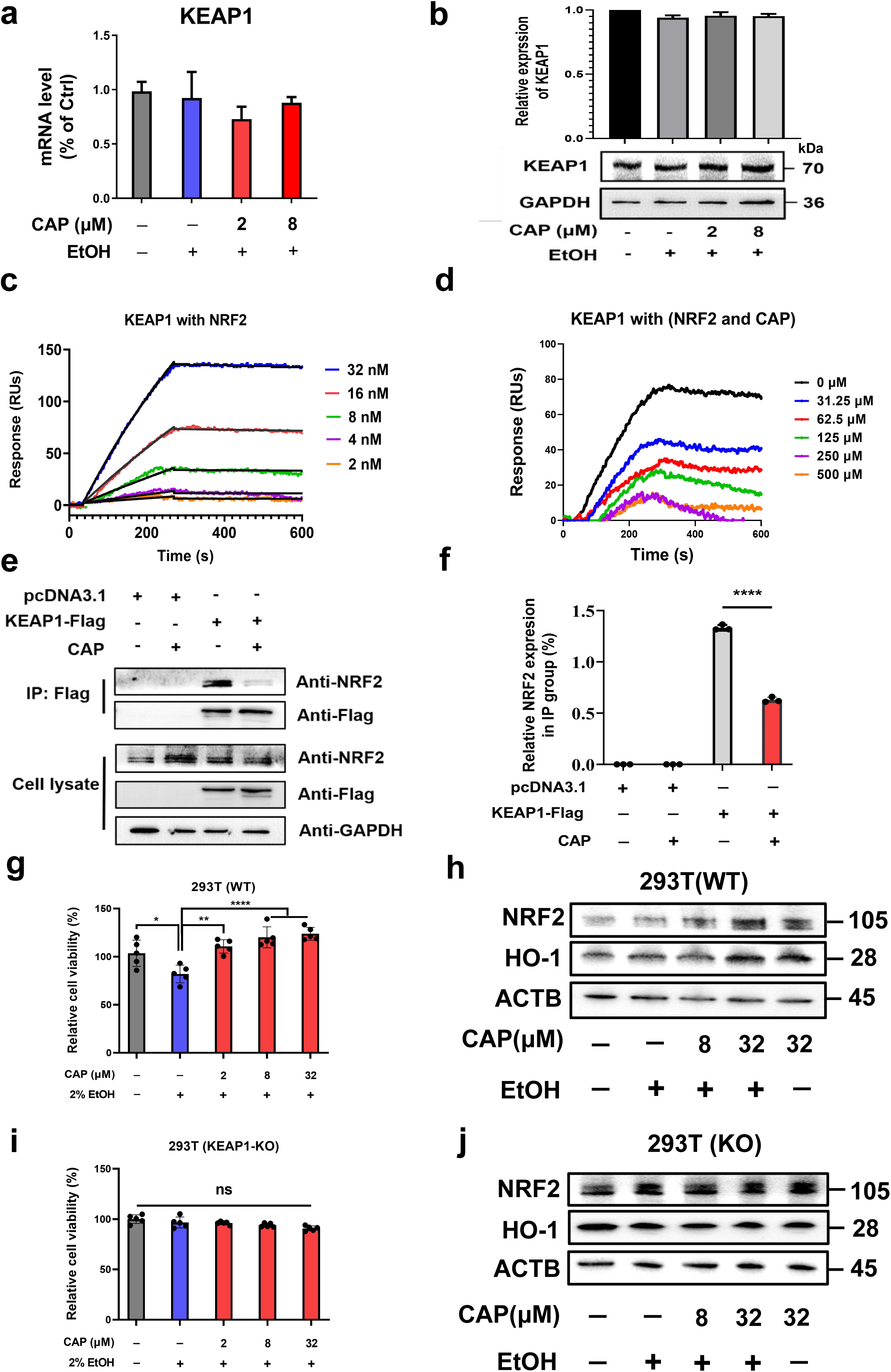
Capsaicin disrupted KEAP1-NRF2 interaction. **(a-b)** Assessment of KEAP1 transcription and expression levels using RT-qPCR and western blot analyses. **(c)** KEAP1-NRF2 interaction was detected with Surface plasmon resonance (SPR) *in vitro*. **(d)** Disruption of KEAP1-NRF2 interaction by CAP as assessed by SPR. **(e)** Disruption of KEAP1-NRF2 interaction by 32 μM CAP as assessed by Co-Immunoprecipitation (Co-IP) in 293T Cells. **(f)** Quantitative analysis of relative NRF2 expression in IP samples using ImageJ software. **(g)** Cell viability assessment of 293T cells treated with CAP and EtOH using CCK-8 assay. **(h)** Western blot analysis of NRF2 and HO-1 expression in 293T cells. **(i)** Cell viability assessment of 293T(KO) cells treated with CAP and EtOH using CCK-8 assay. **(j)** Western blot analysis of NRF2 and HO-1 expression in 293T(KO) cells.

Surface plasmon resonance (SPR) was used to quantitatively evaluate the *in vitro* binding affinity between purified KEAP1 and NRF2. The observed dissociation constant (K_D_) for the KEAP1-NRF2 complex was calculated to be 1.14 ± 0.15 nM **(Figure 3c)**, corroborating earlier reports of a strong association between these two proteins^11^. Upon exposing KEAP1 to increasing concentrations of CAP—ranging from 16 μM to 500 μM—a consistent, dose-dependent decrease in KEAP1-NRF2 binding was noted **(Figure 3d)**. We further confirmed these observations through co-immunoprecipitation (Co-IP) assays. Treatment with 32 μM CAP led to a 52.79% reduction in the endogenous NRF2 associated with KEAP1 in cellular models. **(Figure 3e and 3f)**. Collectively, these results provide the first empirical evidence that CAP effectively impairs the KEAP1-NRF2 interaction.

### 2.7 Knockout of KEAP1 affects the function of CAP

To investigate whether KEAP1 functions as a key target for the antioxidant properties mediated by CAP, KEAP1-knockout (KO) 293T cell lines were generated and validation studies were performed using western blot analysis **(Supplementary Figure 4a and 4b)**. Initially, we observed that CAP significantly activated the NRF2 and HO-1 in wild-type 293T cells, leading to a pronounced increase in cell viability **(Figure 3g and 3h).** This finding was consistent with observations made in GES-1 and UC-MSC cells. However, in KEAP1-KO cells, an innate resilience to oxidative stress was evident, likely due to the constitutive activation of endogenous NRF2 and its downstream effector, HO-1. As a result, the CCK8 assay revealed no significant differences in cell viability between the groups **(Figure 3i)**, corroborating previous studies that have shown KEAP1-KO mice exhibit reduced oxidative stress and inflammatory responses^28^. Importantly, in the KEAP1-KO 293T cell line, CAP displayed a reduced ability to activate NRF2 and HO-1 **(Figure 3j)**. A grayscale quantification of these pivotal proteins was presented in **Supplementary Figure 4d and 4e.** Taken together, our findings strongly support that KEAP1 is a crucial mediator of the antioxidant effects conferred by CAP and plays a pivotal role in amplifying the NRF2-ARE signaling pathway. At the same time, we also partially exclude the possibility that CAP may function through off-target effects involving the TRPV1 receptor or DPP3 with an ETGE motif (**Supplementary Figure 4c and 4f-4i)**.

### 2.8 Capsaicin directly interacts with KEAP1

As previously documented, small molecules categorized as NRF2 agonists, such as Dimethyl fumarate (DMF), curcumin, and sulforaphane, predominantly target and inhibit the KEAP1 function^29^. To probe the interplay between KEAP1 and CAP within a cellular environment, we conducted a cellular thermal shift assay (CETSA). In the absence of CAP, KEAP1 experienced marked degradation within a temperature window of 58°C to 67°C. In contrast, the presence of CAP significantly stabilized KEAP1 against thermal denaturation, resulting in sustained levels of the KEAP1 protein in cells **(Figure 4a)**.

**Figure 4.**
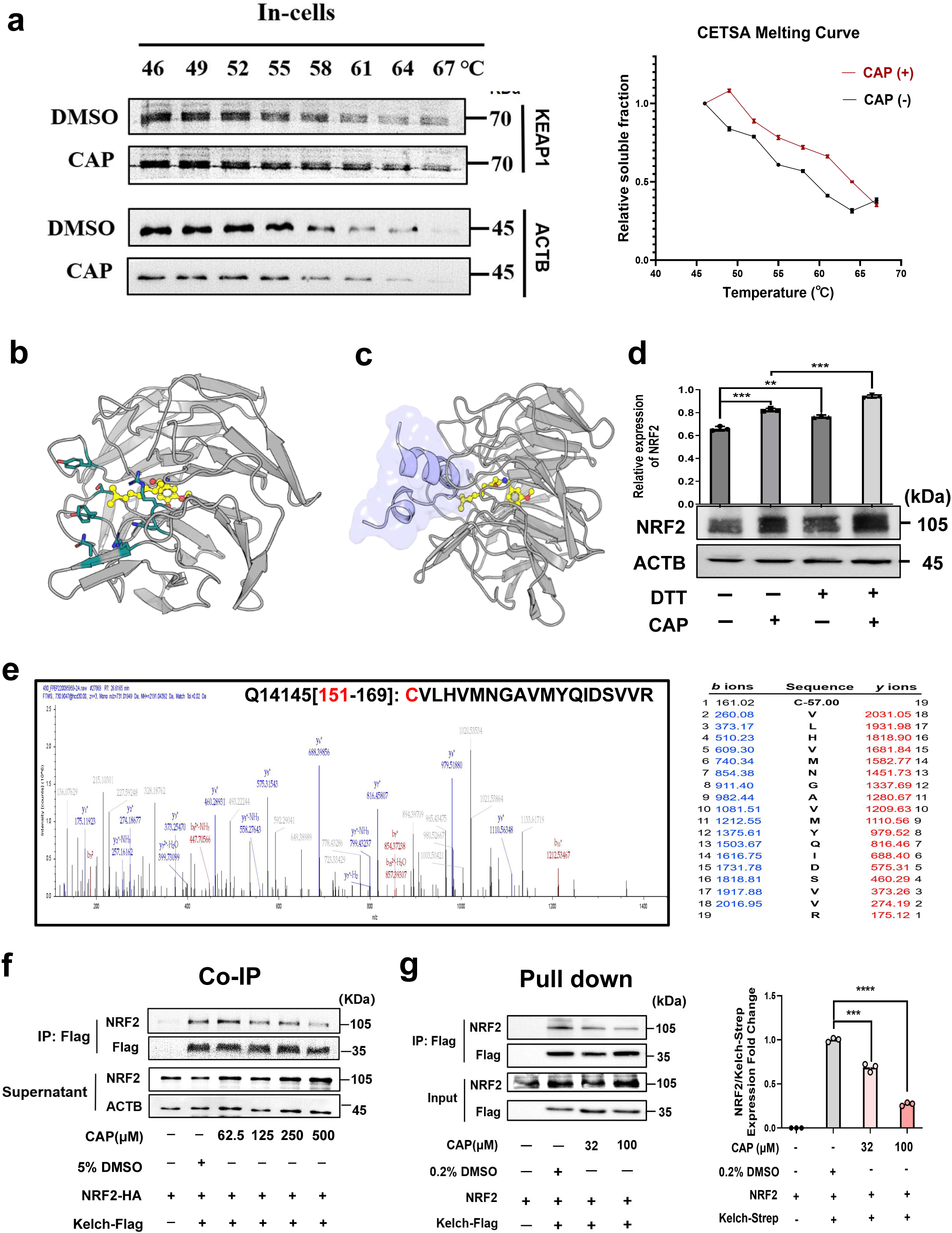
CAP specifically interacts with the Kelch domain of KEAP1. **(a)** Detection of KEAP1-CAP interaction in GES-1 cells using cellular thermal shift assay coupled with western blotting (CETSA-WB) and the melting curve generated from CETSA was analyzed using ImageJ software. The red fold line represents cells treated with CAP, while the black fold line represents cells treated with DMSO as a control. **(b)** Computational docking of CAP molecule to KEAP1 surface pockets. The Keap1 protein is represented in gray, while the CAP molecule is shown in yellow. The seven key amino acids predicted to be crucial for the interaction are highlighted in blue. **(c)** Partial overlap of CAP-binding pocket with KEAP1-NRF2 interface. The KEAP1-NRF2 interaction interface is represented in purple. **(d)** Influence of dithiothreitol (DTT) on NRF2 activation induced by CAP in GES-1 cells. Cells were pre-incubated with 400 μM DTT for 1 hour, followed by a 3-hour incubation with 8 μM CAP. **(e)** Tandem mass spectrometry (MS/MS) of analysis of KEAP1 peptide containing Cys151 following CAP treatment. **(f)** Dose-dependent examination of Kelch-NRF2 interaction in the presence of CAP in 293T cells. Cells were treated with varying concentrations of CAP (0, 62.5, 125, 250, and 500 μM) to assess the impact on the interaction between exogenously purified Kelch protein and NRF2 in the total cell lysate. **(g)** Pull-Down assay demonstrating the direct inhibition of Kelch-NRF2 Interaction by CAP.

To investigate the specific CAP-binding sites on KEAP1, we utilized computational docking and all-atom simulations, to map the positioning of possible CAP-interacting residues within the surface pockets of KEAP1, with a particular focus on the Kelch domain that is responsible for NRF2 interaction **(Figure 4b and Supplementary Figure 5a)**. The most energetically favorable model demonstrated that the vanillyl headgroup of CAP was inserted into the Kelch channel, while its flexible hydrocarbon chain remained at the periphery of the NRF2-Kelch interface. Based on this docking model, we hypothesized that the CAP binding pocket partially overlapped with the KEAP1-NRF2 interacting interface **(Figure 4c)**. Hence, we modeled interactions between NRF2 and Kelch in the presence of CAP using all-atom molecular dynamics simulations. When CAP associated with Kelch, the NRF2-Kelch complex was destabilized **(Supplementary Figure 5b and 5c and Supplementary videos 1 and 2)**. Indeed, for the NRF2-Kelch complex, the average distance between D29 of NRF2 and R415 of Kelch remained approximately 1.20 nm, indicating stable bindings throughout the 100 ns trajectory. For the NRF2-CAP-Kelch complex, however, the average distance fluctuated to 2.87 nm, resulting in the dissociation of NRF2 from the Kelch domain **(Supplementary Figure 5d and 5e)**. Our simulations support the experimental observation that CAP can act as a protein-protein interface inhibitor.

Interestingly, the majority of previously identified KEAP1 inhibitors were known to induce covalent modifications of the cysteine residues of KEAP1, thereby disrupting the interactions between KEAP1-Cul3 and KEAP1-NRF2, and consequently activating the downstream NRF2-ARE signaling pathway^30^. Therefore, we introduced the reducing agent dithiothreitol (DTT) into the cell medium. Existing literature suggests that KEAP1’s active cysteines can be covalently altered by electrophilic agents like CN-2F, and such modifications are generally reversible in the presence of DTT^22^. Additionally, the effectiveness of catechol moieties in NRF2 activation is often diminished by pre-treatment with DTT^31^. However, our results indicated that the CAP-induced elevation of NRF2 expression levels was not inhibited by DTT **(Figure 4d)**.

To further investigate whether CAP also forms covalent bonds with key cysteine residues on the KEAP1 surface, we targeted our scrutiny towards the functionally critical cysteines, such as Cys151, Cys273 and Cys288^32^. Employing tandem mass spectrometry (MS/MS), we assessed the molecular weights of KEAP1 samples following incubation with CAP and found no evidence of CAP-induced modifications **(Figure 4e, and Supplementary Figure 6a-6d)**. These results suggested that although CAP bound directly to KEAP1, it did not do so through covalent interaction.

### 2.9 CAP specifically interacts with the Kelch domain of KEAP1

To investigate the impact of CAP on the binding affinity between the Kelch domain and NRF2, we conducted immunoprecipitation (IP) and Pull-down analysis using purified Kelch-domain-only proteins. The IP results revealed a noticeable reduction in NRF2 proteins bound to the Kelch domain, corresponding with incremental concentrations of CAP. In parallel, the levels of unbound NRF2 in the supernatant progressively increased **(Figure 4f and Supplementary Figure 6e)**. Further substantiating these observations, purified NRF2 proteins were also used. Pull-down assays furnished compelling empirical evidence that CAP effectively disrupted the direct interaction between the Kelch domain and NRF2 **(Figure 4g)**.

### 2.10 CAP binds KEAP1 at allosteric sites

According to the results of molecular docking and dynamic simulation, we purified both the wild-type Kelch domain (WT) and the mutant variant (Mut) of Kelch proteins. The mutant variant involved substitutions at residues Y334A, R380A, N382A, N414A, R415A, Y572A, and S602A (the orthostatic site), which are residues reported to engage NRF2 and traditional Keap1 inhibitors. Bio-Layer Interferometry (BLI) revealed that the binding affinity between CAP and the Kelch domain (WT) was 31.45 μM with R^2^=0.99 **(Figure 5a)**. Intriguingly, the mutated Kelch domain (Mut) exhibited diminished, but still appreciable, affinity for CAP with a K_D_ of 102.4 μM with R^2^=0.98 **(Figure 5b).** Our biophysical examinations suggested the orthostatic NRF2-binding pocket was not entirely responsible for CAP-binding. Indeed, the Kelch (Mut)-Flag protein failed to pull down any NRF2 from cellular lysates, but it still interacted with CAP **(Figure 5c)**. These collective findings suggest that the CAP-binding pocket and the NRF2-binding interface on KEAP1 may be different from each other, implicating CAP as an allosteric modulator of KEAP1. This specificity may arise from the interaction between the vanillyl headgroup of CAP and the main chain atoms of the Kelch domain, which appears to be largely unaffected by NRF2 binding **(Figure 5d)**. The crystal structures depicting the binding sites between previously identified ligands and the Kelch protein (PDB: 4IQK and 5FNQ) were illustrated on the left-hand side of **Figure 5e**. In contrast, the right-hand side revealed our discovery of the binding sites between CAP and the Kelch protein, which were very inconsistent with previous reports. Here, to explore the potential mechanism of allosteric regulation of the Kelch domain with CAP, we performed HDX-MS assays. The addition of CAP led to an increase in the hydrogen-deuterium exchange rate in about three regions of Keap1, L342-L355, D394-G423, and N482-N495, suggesting their conformational changes. It was noteworthy that amino acid residues N414 and R415 are located in the D394-G423 region, which were the key residues for the interaction with NRF2, and CAP binding may lead to their changes, thus CAP may have weakened the interaction between KEAP1 and NRF2 by way of allosteric regulation **(Figure 5f).** These insights suggested that CAP could potentially represent a novel class of NRF2 agonists.

**Figure 5.**
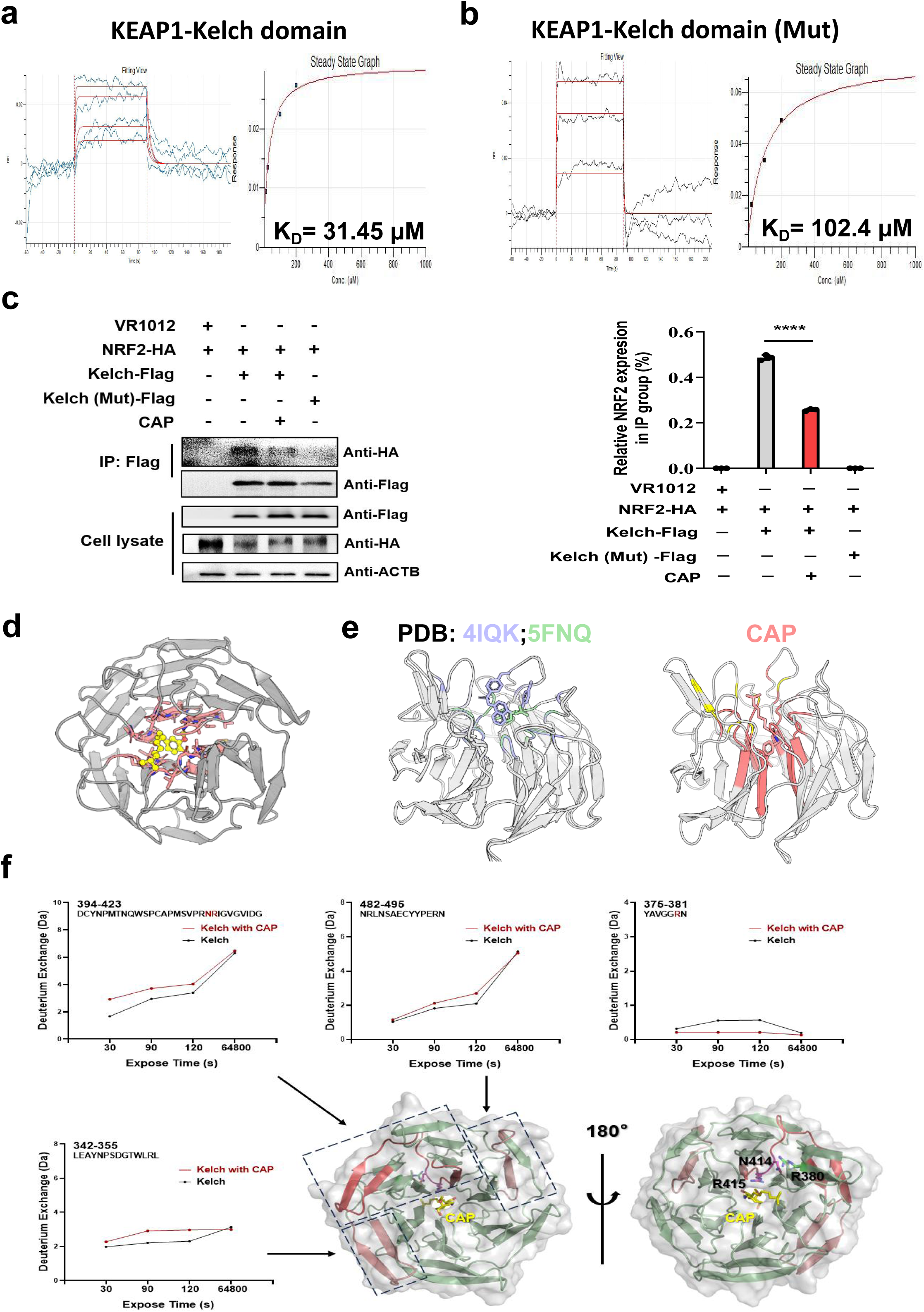
Mutation of KEAP1 affects the function of CAP. **(a)** *In vitro* detection of KEAP1-Kelch domain with CAP using BLI. **(b)** *In vitro* detection of KEAP1-Mutant Kelch domain (Y334A, R380A, N382A, N414A, R415A, Y572A and S602A) with CAP using BLI. **(c)** Co-IP assay to assess the interaction between mutant Kelch and NRF2, and the Impact of 32 μM CAP on Kelch-NRF2 binding in 293T cells. **(d)** Docking analysis reveals encirclement of CAP’s vanillyl headgroup by main chain atoms of the Kelch domain. **(e)** Left side:existing binding sites of two common ligands with Kelch (PDB:4IQK and 5FNQ); Right side:newly discovered binding sites of CAP with Kelch. **(f)** CAP allosterically regulated the conformation of Kelch by HDX-MS. Peptides with increased deuterium uptake ratio after CAP treatment are highlighted in red.

### 2.11 Preparation and characterization of IR-HSA@CAP

Encouraged by the discovery of CAP as a novel agonist of NRF2, we developed a convenient nano-drug delivery platform for the efficient utilization of CAP *in vivo*. We opted for Human Serum Albumin (HSA) as the encapsulating material, given its FDA-approved status for biocompatibility. HSA has also been documented to offer protective effects on the gastric mucosa^33^. As a fluorescent marker, we employed IRDye800, a widely utilized near-infrared fluorescent dye in biomedical research for applications such as protein tagging and imaging applications. Utilizing these components, we successfully formulated CAP-encapsulated IRDye800-HSA nanoparticles, referred to as IR-HSA@CAP **(Figure 6a)**. The labeling ratio stood at 1.48 (IRDye800: HSA) and achieved a coupling efficiency of 93.29% **(Figure 6b)**, and the encapsulation was notably high as calculated^34^.

**Figure 6.**
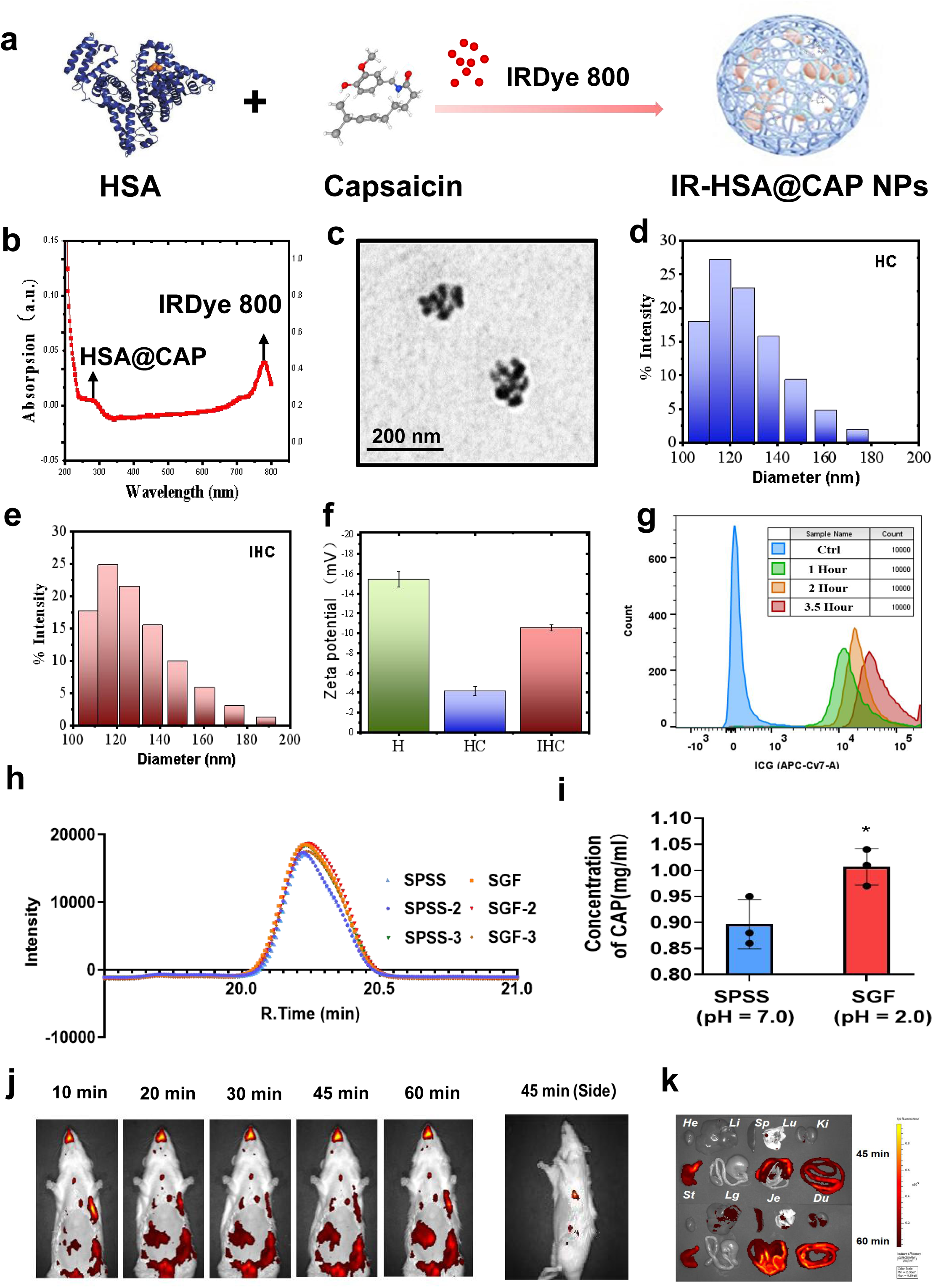
Preparation and characterization of IR-HSA@CAP NPs. **(a)** Schematic illustration of the synthesis process for IR-HSA@CAP nanoparticles (NPs). **(b)** Determination of IRDye800 binding with HSA by UV-Vis spectra. **(c)** The morphology of IR-HSA@CAP NPs was investigated by transmission electron microscope. Scale bar, 0.5 μm. **(d-e)** Particle size comparison of HSA@CAP and IR-HSA@CAP NPs. **(f)** Zeta-Potential analysis of nanomaterials. The zeta-potential values of the different materials were measured: H representing HSA, HC representing HSA@CAP NPs, and IHC representing IR-HSA@CAP NPs. **(g)** FCM analysis demonstrates endocytosis of IR-HSA@CAP NPs by GES-1 cells. **(h-i)** Detection of CAP release in simulated gastric fluid (SGF) and stroke-physiological saline solution (SPSS) Using high-performance liquid chromatography (HPLC). **(j)** *In vivo* imaging reveals the localization of IR-HSA@CAP NPs. **(k)** Observing the distribution of IR-HSA@CAP NPs in major organs of rats.

The structural attributes of the IR-HSA@CAP nanoparticles were characterized using transmission electron microscopy (TEM), the results of which were illustrated in **Figure 6c**. Notably, the size distributions of HSA@CAP and IR-HSA@CAP nanoparticles were remarkably similar. The former exhibited a mean particle size of 139.27 ± 7.49 nm **(Figure 6d)**, while the latter registered at approximately 160.97 ± 14.78 nm **(Figure 6e)**. In terms of Zeta-potential, the HSA@CAP nanoparticles displayed an increased to -4.16 ± 0.78 mV post-formation, as compared to native HSA, which had a potential of -15.45 ± 1.40 mV. Interestingly, this Zeta-potential returned to a value of -10.55 ± 0.50 mV after incorporating IRDye800 in the IR-HSA@CAP **(Figure 6f).**

Importantly, no discernible decline in cell viability was observed over a 24-hour period in both GES-1 and RAW264.7, underscoring the non-cytotoxicity of the nanoparticles **(Supplementary Figure 7a and 7b)**. Furthermore, intracellular fluorescence intensity escalated continuously with time, reaching a point where nearly all cells exhibited IR-Dye800 positivity within one hour **(Figure 6g)**. These findings strongly implied that the IR-HSA@CAP nanoparticles were rapidly internalized by cells and show excellent biocompatibility.

To precisely quantify the CAP content in IR-HSA@CAP nanoparticles, we established a standard curve for CAP using high-performance liquid chromatography (HPLC) at a wavelength of 282 nm **(Supplementary Figure 7c)**. To evaluate the stability and release profile of the nanoparticles in gastrointestinal conditions, we examined the release kinetics of CAP in normal saline (SPSS, pH=7.0) and simulated gastric fluid (SGF, pH=2.0). Initial tests revealed that virtually all encapsulated CAP could be rapidly released in the SGF **(Supplementary Figure 7d and 7e)**. In subsequent animal trials calibrated to the actual concentrations of CAP, we found that the CAP concentration in SGF swiftly attained a theoretical value of 1 mg/ml within 30 minutes, while remaining essentially constant in SPSS conditions **(Figure 6h and 6i)**. These findings collectively suggest that IR-HSA@CAP nanoparticles were capable of achieving rapid release of CAP in the gastric milieu to a notable extent.

### 2.12 IR-HSA@CAP NPs activated the Nrf2-ARE signaling pathway *in vivo*

To investigate the protective effects of IR-HSA@CAP nanoparticles against EtOH-induced gastric mucosal injury *in vivo*, Sprague-Dawley rats were randomly assigned to eight different experimental groups. Rebamipide was selected as the positive control due to its well-established ability to protect gastric mucosa^35^.

The *in vivo* experimental protocol was conducted as follows: To ensure gastric emptying, rats were subjected to a 24-hour fasting period. Subsequently, they were administered SPSS, CAP, or the designated treatment in accordance with their group allocation. After a 45-minute pretreatment phase, the rats were exposed to 75% EtOH for an additional 1.5 hours. Following this, the animals were humanely euthanized, and the gastric tissue samples were collected for subsequent experiments and analyses. Precise dosages administered were as follows: CAP at 1 mg/kg, HSA at 2 mg/kg, and Rebamipide at 100 mg/kg. The HSA@CAP nanoparticles used were prepared to contain 1 mg/kg of CAP and 2 mg/kg of HSA, as quantified by HPLC three times.

To elucidate the spatiotemporal distribution of CAP within the rat model, imaging was performed at various time intervals following administration. The data revealed that CAP was predominantly absorbed by the gastrointestinal tract, exhibiting significant localization in the stomach **(Figure 6j and 6k)**.

The administration of either CAP or IR-HSA@CAP alone did not result in detectable damage to the gastric tissues, in contrast to the control group. However, significant hemorrhaging and erosion were evident in gastric tissues exposed to EtOH **(Figure 7a, left)**. Utilizing the Guth scoring system, we quantitatively assessed the Gastric Mucosal Ulcer Injury (UI) index. The average UI index in the EtOH group increased to 36.0, indicating acute tissue damage. On the other hand, significant reductions in the UI index were observed in both the CAP and IR-HSA@CAP pretreatment groups, registering at 7.0 and 5.3 respectively. These levels were similar to the Rebamipide group, which had a UI index of 9 **(Figure 7a, right)**. These findings align well with our expectations. Notably, the effective dose of CAP (1 mg/kg) was substantially lower than that of Rebamipide (100 mg/kg), underscoring the potential advantages of CAP. Additionally, while the HSA-pretreated group did exhibit some attenuation of gastric mucosal injury, the reduction was not statistically significant (UI=17.7). This suggested that CAP was the principal active component responsible for the observed effects in the IR-HSA@CAP nanoparticles.

**Figure 7.**
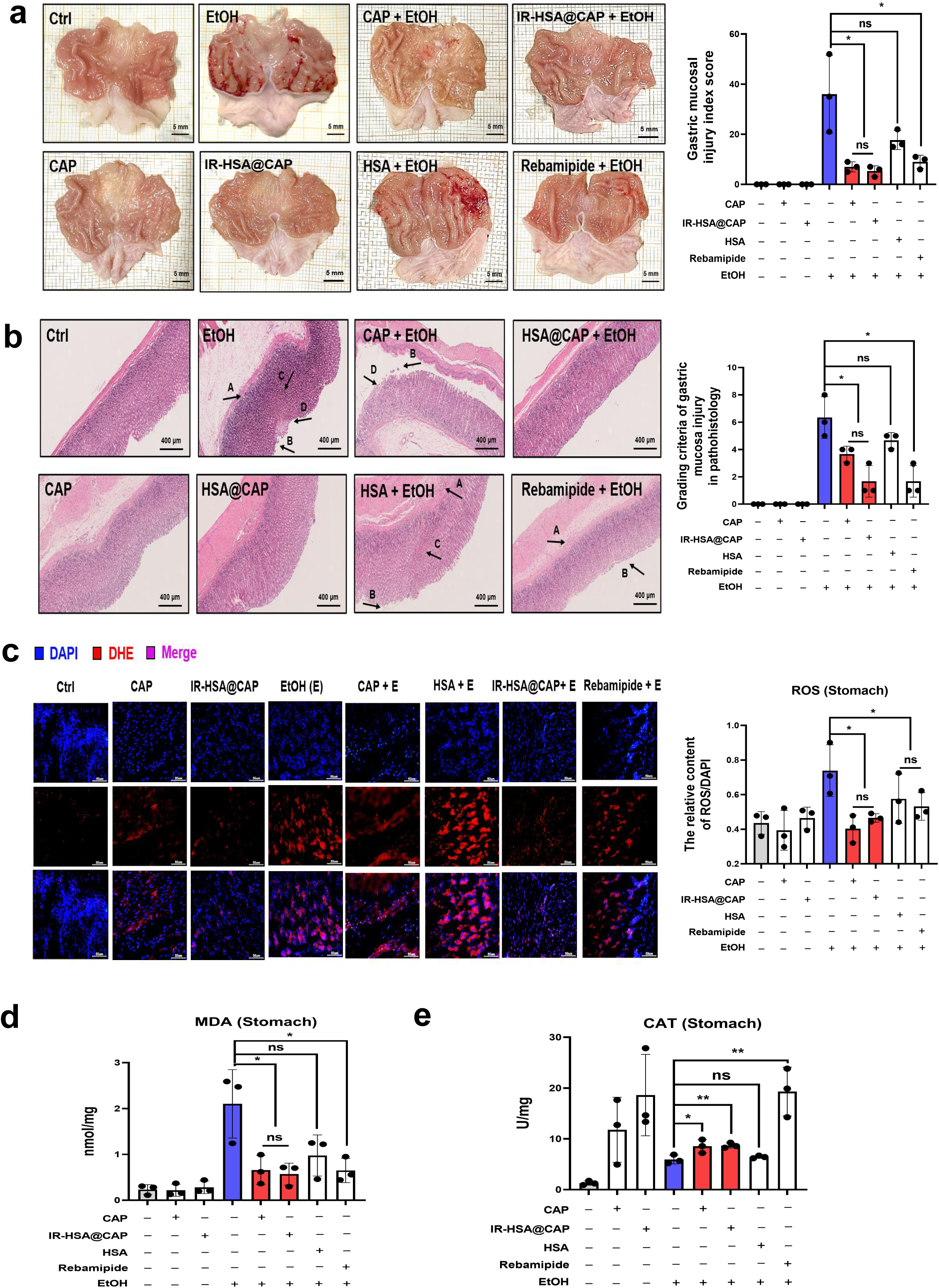
CAP activated NRF2-ARE pathway *in vivo*. **(a)** Impact of CAP (1 mg/kg) on histopathology of EtOH-induced gastric mucosal injury. Rebamipide (100 mg/kg) was used as a positive control. Tissue sections were stained and evaluated for gastric mucosal ulcer injury using the Guth scoring system. Representative images were shown with a scale bar of 5 mm. The ulcer injury (UI) index was calculated. **(b)** Histological examination of rat gastric mucosa using H&E staining. Scale bar represents 400 μm. A: Inflammatory cell infiltration; B: Epithelial exfoliation; C: Glandular disorder; D: Gastric edema. Quantitative analysis was conducted using the Masuda scoring system. **(c)** Visualization of ROS in rat gastric tissue under various treatments using DHE staining and inverted fluorescence microscopy. Scale bar represents 50 μm. ROS levels were quantified using Image Pro Plus 6.0 software. **(d)** MDA levels in gastric tissues across different treatment groups. **(e)** Assessment of catalase (CAT) activity in gastric tissues.

Subsequently, we quantified the extent of pathological damage using H&E staining coupled with the Masuda scoring system, wherein a higher index reflects more severe tissue damage. The pathological damage index registered at 6.3 in the EtOH-only group, while the HSA-treated group showed a slightly lower index at 4.7. The CAP-treated group further reduced this index to 3.7. Remarkably, the employment of either IR-HSA@CAP nanoparticles or Rebamipide led to a substantial decline in pathological damage, with both treatments registering an identical index of 1.7 **(Figure 7b)**. These results demonstrate that the therapeutic efficacy of IR-HSA@CAP nanoparticles surpassed that of either CAP or HSA administered individually. Specifically, the reduction in tissue damage was 1.77-fold greater than that achieved with CAP alone and 2.88-fold greater compared to HSA alone.

To evaluate the capability of CAP to inhibit the EtOH-induced augmentation of ROS levels in gastric tissues *in vivo*, we conducted dihydroethidium (DHE) staining on cryo-sectioned gastric specimens. Pre-treatment with either CAP or IR-HSA@CAP nanoparticles substantially mitigated ROS production, with reductions of 45.42% and 36.99% respectively, compared to the EtOH-only group **(Figure 7c)**. Concurrently, MDA assays underscored that IR-HSA@CAP nanoparticles efficaciously curtailed lipid peroxidation levels in the gastric tissues by an approximate 72.82% **(Figure 7d)**. Additionally, CAT activity in the tissues was enhanced by 31.85% following treatment with IR-HSA@CAP nanoparticles (**Figure 7e)**.

Furthermore, we scrutinized the modulatory effects of IR-HSA@CAP on the Nrf2-ARE signaling pathway. Our findings, substantiated through immunohistochemistry **(Figure 8a-8c)** and western blot analyses **(Supplementary Figure 8j-8m)**, revealed that pre-treatment with IR-HSA@CAP led to a significant upregulation of Nrf2 expression. Concurrently, IR-HSA@CAP pre-treatment also stimulated elevated expression levels of HO-1 and Trx. These collective observations suggest that IR-HSA@CAP was capable of orchestrating the Nrf2-ARE signaling axis, thereby providing a rapid and effective modulatory mechanism for oxidative stress and homeostatic regulation. To delve deeper into the anti-inflammatory potential of our treatment strategy, we analyzed homogenized gastric tissue samples for the expression profiles of various chemokines and inflammatory mediators. Intriguingly, pre-treatment with either CAP or IR-HSA@CAP elicited a marked suppression in the expression levels of pro-inflammatory cytokines such as IL-1β, TNF-α, IL-6, and CXCL1, while simultaneously augmenting the expression of the anti-inflammatory cytokine IL-10 **(Figure 8d).** Not surprisingly, IR-HSA@CAP exhibited a stronger anti-inflammatory effect. The novel mechanism of capsaicin nanoparticles in preventing EtOH-induced gastric mucosal injury is shown in **Figure 8e**.

**Figure 8.**
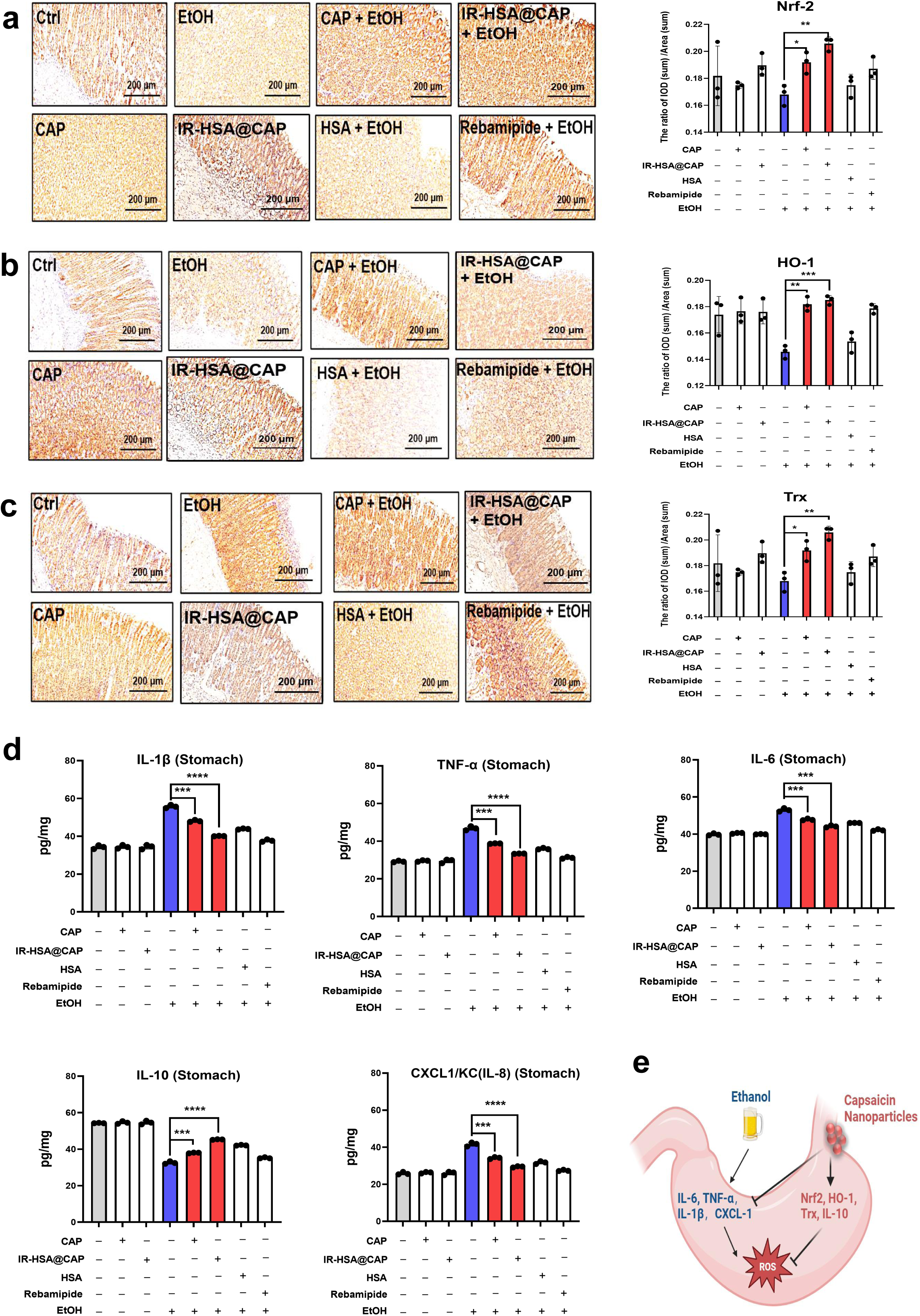
CAP also suppressed inflammation *in vivo*. **(a-c)** Immunohistochemical detection of antioxidant proteins and presentation of representative images and statistical analysis results. (a) Nrf2, (b) HO-1, and (c) Trx protein expression levels were assessed. **(d)** ELISA measurement of IL-1β, TNF-α, IL-6, CXCL1/KC (IL-8), and IL-10 Levels in gastric tissues. Data were presented as mean ± SD. Significance levels were indicated as follows: ns, not significant; *p < 0.05; **p < 0.01; ***p < 0.001; ****p < 0.0001 compared to the EtOH-only group (marked in blue). **(e)** Mechanism diagram of capsaicin alleviating gastric mucosal injury caused by ethanol in rats.

Finally, we assessed the biosafety of IR-HSA@CAP in rats through continuous oral administration over a one-week period **(Supplementary Figure 8a)**. No significant alterations were observed in the organ indices for either the liver or spleen (**Supplementary Figure 8b and 8c)**. Comprehensive blood biochemical analyses further revealed that varying concentrations of IR-HSA@CAP exerted no deleterious impact on hepatic function, as indicated by levels of ALT, AST, and T-Bil **(Supplementary Figure 8d-f)**, or on renal function, as evidenced by measurements of BUN, CRE, and UA **(Supplementary Figure 8g-i)**. These collective findings underscore the high biosafety profile of the IR-HSA@CAP nanoparticles.

### 2.13 Protective effect of IR-HSA@CAP NPs was partially eliminated in Nfe2l2-KO mice

A direct proof that IR-HSA@CAP NPs act through activation of Nrf2 in protecting gastric mucosa against alcohol toxicity could be well conducted in commercially available Nfe2l2-knockout mice. Therefore, we conducted experiments in normal C57BL/6 mice and Nfe2l2 knockout mice.

The phenotype and H&E staining were shown in **Figure 9a and Figure 9b**. In normal mice, EtOH can still cause significant gastric mucosal damage, and our drug also had a beneficial effect on mice. In the Nfe2l2-knockout group, we found that these mice showed more sensitivity to EtOH. The gastric injury of model group without Nrf2 expression was more obvious than normal mice given the same amount of EtOH, because we found significant gastrointestinal bleeding **(Figure 9c, a representative photograph)**. This suggested that Nrf2 may be important for mice to cope with gastric mucosal damage caused by EtOH.

**Figure 9.**
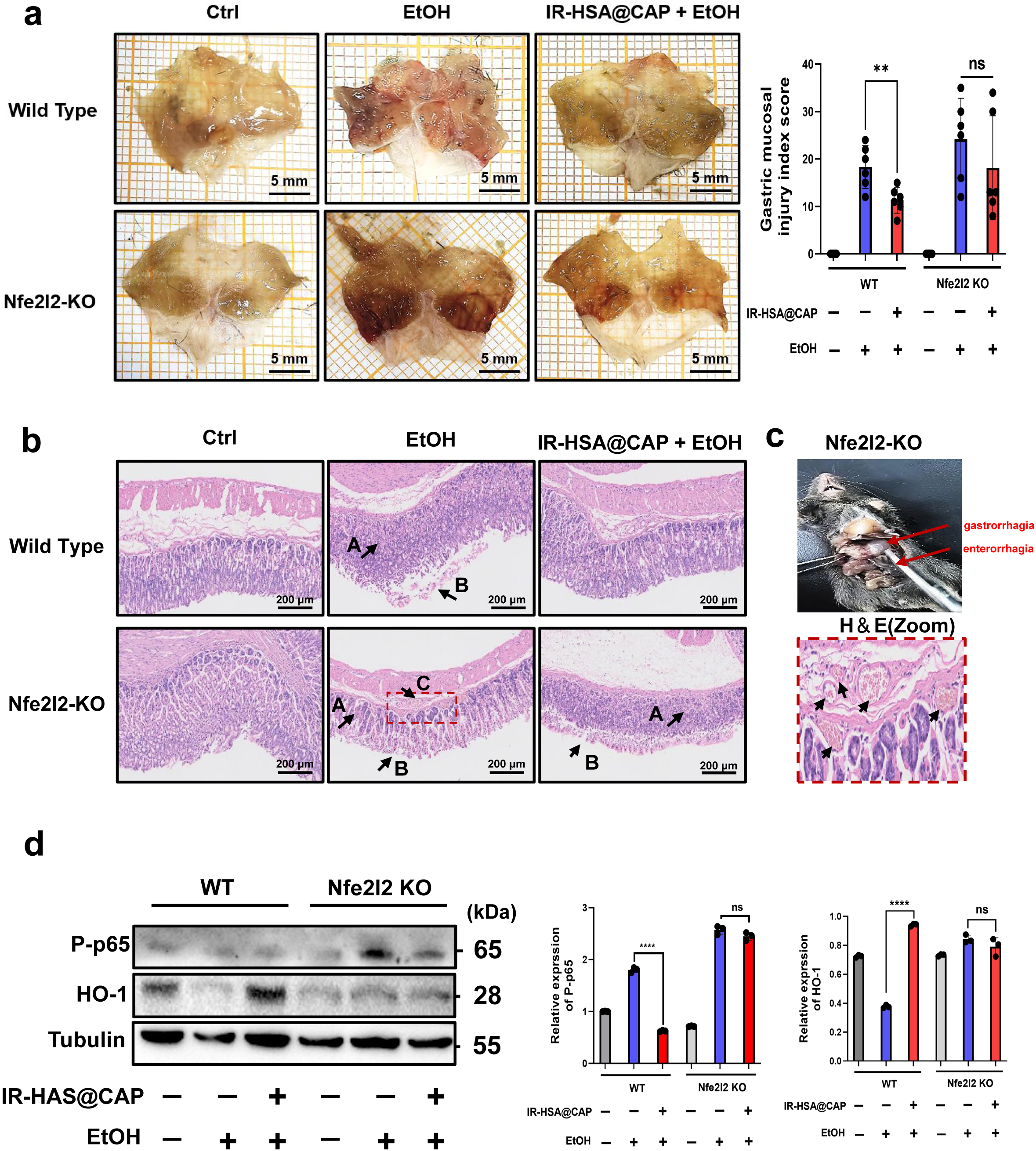
Protective effect of CAP was partially eliminated in Nfe2l2-KO mice. **(a)** Tissue sections were evaluated for gastric mucosal ulcer injury using the Guth scoring system. Representative images were shown with a scale bar of 5 mm. The ulcer injury (UI) index was calculated. **(b)** Histological examination of mice gastric mucosa using H&E staining. Scale bar represents 200 μm. A: Glandular disorder; B: Epithelial exfoliation; C: Inflammatory cell and erythrocyte infiltration. **(c)** Gastrointestinal bleeding and H&E staining (Zoom) in a representative Nfe2l2-KO mouse treated with EtOH. **(d)** Western blotting to detect the key proteins of P-p65 and HO-1.

At the same time, the gastric injury after drug treatment was only partially reduced in Nfe2l2-deficient mice, indicating that the IR-HSA@CAP NPs played a key role by activating Nrf2 *in vivo*. Here, we also selected two proteins related to inflammation (phosphorylated p65) and antioxidant (HO-1, one of the most critical downstream genes of NRF2) for further western blotting experiments. The results showed that CAP could play an antioxidant and anti-inflammatory role in normal mice, while the related proteins in knockout mice were not significantly different from those in the model group **(Figure 9d)**. Additional experiments were conducted on rats, as shown in **Figure 7 and Figure 8**. These results were consistent with our expectations that IR-HSA@CAP NPs act through activation of Nrf2 in protecting gastric mucosa against alcohol toxicity.

## 3 Discussion

CAP, a natural small molecule derived from chili peppers, has been identified and verified to modulate the KEAP1-NRF2-ARE pathway in our study. This signaling pathway plays a key role in several cancerous and non-cancerous diseases^36–40^. Therefore, it is a promising therapeutic target for reducing free radical damage associated with numerous conditions, including Alzheimer’s disease^41^, chronic kidney disease^42^, cardiovascular disease^43^, cancer and aging^44^.

Over the past few decades, the scientific community has tirelessly sought molecules capable of activating NRF2. Hu et al. reported the first small-molecule inhibitor of the Keap1-Nrf2 PPI from a high-throughput screen (HTS) in 2013^45^. Davies and co-workers began with a crystallographic screen of approximately 330 fragments and developed KI-696 as a nanomolar KEAP1-NRF2 PPI inhibitor^46,47^. A spectrum of NRF2 agonists, both natural and synthetic, covalent and non-covalent, have been unveiled. However, Kim T. Tran et al. reported that many compounds are difficult to synthesize and the binding activity of half of the compounds cannot be demonstrated convincingly^48^. Practical application faces two primary challenges. First and foremost, covalent modifications to KEAP1 pose a noteworthy concern regarding “off-target” effects owing to their electrophilic properties. These effects have the potential to impact various other cellular proteins, impeding their commercial success^29^. Furthermore, unregulated activation of NRF2 can inadvertently foster cancer progression by protecting malignant cells and promoting their proliferation, metastatic potential, and resistance to therapeutic interventions^49^.

Recent advancements have been geared towards the development of direct inhibitors for the KEAP1-NRF2 interaction, drawing significant attention from the scientific community^50^. Interestingly, our research unequivocally identifies the non-covalent interaction between CAP and KEAP1. Dimethyl fumarate (DMF), a KEAP1-dependent NRF2 agonist, has progressed positively in phase III clinical trials. The recognition that DMF’s minimal side effects stem from its dual mode of action—both covalently modifying reactive cysteines and non-covalently activating NRF2 via the Kelch domain of KEAP1—has marked a pivotal milestone in the development of drugs targeting KEAP1^29^. We found that at lower concentrations, CAP effectively bound with the Kelch domain **(Figure 5a and 5c)** and activated NRF2, subsequently promoting the expression of downstream antioxidant genes **(Figure 1i-1m)**. *In vivo* studies, we observed almost no adverse reactions. Additionally, Merritt, J.C. et al. reported that CAP at high concentrations possesses potent anti-cancer properties^51^. This aligns with our CCK8 experimental findings that heightened CAP concentrations curtailed cell viability **(Supplementary Figure 1c and 1d)**. The duality of pharmacological impact of CAP remains a topic of discussion. The interplay and mechanisms between CAP’s varying modes in both regular and cancerous cells, which underlie its antioxidant and anticancer actions, are intricate and warrant further exploration.

Notably, to date, no natural product has been confirmed as a direct inhibitor of the KEAP1-NRF2 interaction. In our findings, the dissociation constant (K_D_) between CAP and Kelch was reported as 31.45 μM **(Figure 5a)**, a relatively weak association compared with ML334 (K_D_=1 μM) and some synthetically designed peptides^52^. However, the KEAP1-NRF2 interaction was significantly inhibited by CAP **(Figure 3d and 3e)**. Of particular significance and surprise, our investigation revealed that CAP and Kelch establish interactions at allosteric sites. Furthermore, HDX-MS was first used to identify the potential allosteric regulatory site of KEAP1 by natural small molecule. In particular, the deuterium exchange of the new peptide (D394-G423) we found was significantly increased, and it was also observed in a newly screened natural small molecule and Kelch (data not shown). Our comprehensive analysis of the CAP-KEAP1 binding dynamics, particularly the unique mode of interaction with Kelch, offers valuable insights for the development of innovative NRF2 activators.

Nonetheless, several concerns warrant further examination. The impact of CAP on the gastric mucosa, as evidenced in animal studies, is contingent upon its dosage and duration of administration. A low dose of CAP applied short-term can mitigate indomethacin-induced gastric microhemorrhage. Conversely, a high dose applied prophylactically over an extended period does not confer this benefit ^53, 54, 55^. It was similarly observed in our studies, where the effective concentration window for natural CAP is a somewhat narrow. For individuals with pre-existing gastric conditions, the epithelial cells of the gastric mucosa might exhibit heightened sensitivity to CAP. This not only diminishes its protective capability but might also exacerbate ulcer severity. Concurrently, the IR-HSA@CAP NPs we employed demonstrated prompt gastric release *in vivo*, underscoring the superior attributes of nanomedicine. This could amplify the antioxidant and anti-inflammatory potency of CAP, thereby resulting in enhanced therapeutic outcomes. Yet challenges persist, particularly the inability to achieve a treatment that targets the stomach exclusively. A limitation of this study is the use of only three rats per group for the *in vivo* experiments. Considering ongoing research into nanomaterials for CAP, there is an opportunity to further refine CAP’s delivery strategy from a pharmacological standpoint, and in future related studies, we will strive to increase the sample size to enhance the robustness of our findings.

In summary, the present study provides evidence that CAP effectively mitigates oxidative stress through the activation of NRF2 and its non-covalent binding to KEAP1’s Kelch domain. The unique binding site between CAP and Kelch distinguishes from the commonly reported small molecule inhibitors, hinting at its potential as a novel class of NRF2 agonists. Moving forward, the development of derivatives inspired by CAP holds promise as a compelling avenue for the creation of therapeutic agents aimed at attenuating oxidative damage.

## 4 Materials and methods

### Reagents and chemicals

Capsaicin (CAP) used *in vitro* was purchased from Solarbio Life Science (CAS:404-86-4, Cat No. IC0060). CAP used *in vivo* was purchased from MCE (Cat No. HY-10448). The anhydrous ethanol (EtOH) was purchased from Tianjin Jiangtian Chemical Technology Co. LTD in China (CAS:64-17-5, Cat No.268). Primary antibodies of anti-HA-Tag (Cat No. AE008), anti-Strep II-Tag (Cat No. AE066) and anti-MYC-Tag (Cat No. AE010) were purchased from ABclonal Technology. The primary antibody to anti-Flag (Cat No. 80010-1-RR), anti-Actin (catalog no. 66009-1-Ig), anti-GAPDH (Cat No. 60004-1-Ig), anti-NRF2 (Cat No. 16396-1-AP) and anti-NRF2 in revised version (Cell Signaling Technology, Cat. No. 12721), anti-GSS (Cat No. 15712-1-AP), anti-HMOX1(Cat No. 10701-1-AP) anti-TXN (Cat No. 14999-1-AP) and anti-KEAP1(Cat No. 30041-1-AP) were purchased from Proteintech in China. Anti-HA affinity matrix (Cat No. 11815016001) was purchased from Roche. Anti-Flag affinity matrix (catalog no. A2220) and albumin from human serum (HSA, Cat No. A3782) was purchased from Sigma. The FITC Goat anti-Rabbit IgG (H+L) was purchased from SIMUBIOTECH (Cat No. S2003). Reactive Oxygen Species Assay Kit (Cat No. CA1410), DAPI (Cat No. H-1200) and the CCK-8 cell proliferation and cytotoxicity assay kit (Cat No. CA1210) were acquired from Solarbio Life Science. The enhanced chemiluminescence (ECL) reagent (Cat No. SQ201) was obtained from EpiZyme. HSA Cycloheximide (CHX, Cat No. HY-12320), DTT (Cat No. HY-15917) and Bortezomib (PS-341, Cat No. HY-10227) were purchased from MCE. Human KEAP-1 Protein was bought from Sino Biological (Cat No.11981-HNCB). Recombinant human NRF2 protein was bought from Abcam (Cat No. ab132356). IRDye® 800CW was bought from *LI-COR.* Nfe2l2-KO mice (Cat. NO. NM-KO-190433) were purchased from Shanghai Model Organisms Center, Inc.

### Cell lines and cell culture

Human gastric mucosal epithelial cells (GES-1), HEK293T human embryonic kidney 293 cells (293T) and Human umbilical cord mesenchymal stem cells (UC-MSC) were obtained from the American Type Culture Collection (ATCC) and cultured at 37°C in 5% CO_2_. KEAP1 Knockout 293T Cell lines (293T-KO) were purchased from ABclonal (RM02350). 293T and 293T-KO were maintained in Dulbecco’s modified Eagle’s medium (DMEM) supplemented with 10% fetal bovine serum (FBS) and 1% penicillin-streptomycin (100 IU/mL). GES-1 and UC-MSC were maintained in Roswell Park Memorial Institute 1640 medium (RPMI-1640), supplemented with 10% fetal bovine serum (FBS) and 1% penicillin-streptomycin (100 IU/mL), RAW 264.7 also needs an additional 1%Glutamax. The cell passage ratio was 1:2-1:4, every 2-3 days.

### Plasmid construction

The plasmids of pCDNA3.1-KEAP1-FLAG (Cat No. P8893) and pCMV-NFE2L2 (human)-HA-Neo (Cat No. P45110) were purchased from the platform of MiaoLingPlasmid (http://www.miaolingbio.com/). In order to better express NRF2 in 293T, we inserted the NFE2L2 (human)-HA into EcoRV/Xba□ of VR1012 through the pEASY®-Basic Seamless Cloning and Assembly Kit purchased from Transgen Biotech (Cat No. CU201-02). The plasmid of Ub-K48-Myc and the no-load plasmids of pcDNA3.1 and VR1012 are available in our own laboratory. The Kelch domain of KEAP1(Lys312-Cys624)^56^ was downloaded and then the optimized (for Escherichia coli; Homo sapiens) sequence was cloned into the vector of pET-28a (+) by adding 5’ (NdeI) and 3’ (XhoI), so the plasmid of kelch in pET-28a(+) was constructed from the company of Azenta. At the same time, a Kelch plasmid with mutation at 7 sites (R415A, N382A, R380A, N414A, S602A, Y334A and Y572A) was designed, which was also synthesized directly using the same method described above.

### CCK-8 cell viability and cytotoxicity assay

GES-1 cells cultured in 96-well plates were used for CCK8 experiments. After the cells had been cultured for more than 24 hours and spread all over the bottom of the plate (10^4^ cells/ml), the experiments were carried out. The cells were washed twice with PBS, low, medium and high concentrations of ethanol (EtOH) were added into the medium of 1640 (0.5%, 3.5% and 5% v/v) and cultured for 1.5 h at 37°C in 5% CO_2_. After the 1.5 h, carefully discard the supernatant before slowly adding the well diluted CCK8 solution, and the absorbance at 450 nm was measured using a microplate reader according to the manufacturer’s instructions. To identify the effective concentration of CAP, CAP dissolved in DMSO (original concentration of 50 mM) was diluted to the specified concentration in 1640 medium(2, 4, 8, 16 and 32 μM), mixed well and added to the cells for 2.5 h. and then added the corresponding concentration of EtOH directly for another 1.5 h. After a total of 4 hours of treatment, CCK8 cell viability was measured according to the above method. UC-MSC cells was treated like GES-1, using 200 μM H_2_O_2_ to replace EtOH. What’s more, in order to judge the safety of nanomaterials, HSA@CAP NPs including HSA (0.04 mg/ml) and CAP (0.02 mg/ml) were added into GES-1 and RAW264.7 cells for 24 hours and then the cytotoxicity assay was conducted same as the tests for cell viability.

### Effect of CAP and EtOH on the morphology of GES-1

GES-1 cells with good condition were cultured in 24-well plates at a concentration of 5 × 10^5^/well. Then, 5% EtOH was added in the EtOH group and the cells pre-added with 2 μM or 8 μM of CAP were termed as the “2 μM CAP + EtOH” group and the“8 μM CAP + EtOH” group. Cell morphology of each group was observed under the fluorescence microscope using only the bright field. In subsequent cell experiments, we also adopted the same modeling method and set such four groups, hereinafter referred to as the model construction of this experiment.

### Flow cytometry of apoptosis

GES-1 cells cultured in 6-well plates were used for detecting apoptosis. Annexin V-FITC Apoptosis Detection Kit (Solarbio Life Science) was used to determine the cell apoptosis rate according to the manufacturer’s instructions. In 6-well plates, GES-1 cells were seeded. The treatment of the cells was consistent with that of the morphological examination. Then, we washed them with cold PBS three times and used a cell scraper to carefully collect cells in each group before staining with Annexin V-FITC and PI at room temperature (RT) for 15 min. Finally, flow cytometry was used to detect apoptotic cells using a FACS flow cytometer (BD Biosciences, USA). The results were analyzed by FlowJo software (version 10.8.1, TreeStar, USA).

### ROS detection

For testing the intracellular ROS, 10^6^ GES-1 cells were cultured on a 12-well plate and placed in an incubator at 37 °C under 5% CO_2_. After CAP pretreatment for 2.5 h and EtOH incubation for 1.5 h, the intracellular ROS level was measured. 1 ml of DCFH-DA (10 μM) was added and incubated in the dark at 37 °C for 30 min. Then, we collected cells and washed them with PBS three times, and finally, the ROS activity was detected by inverted fluorescence microscope and flow cytometry. For the detection of ROS in gastric tissue, five micrometers cryosections of frozen gastric tissues were stained with 10 µM Dihydroethidium (DHE) for 30 min in the dark at RT. Pictures were taken at 200-fold magnification and quantification by Ipp6. 0.

### Measurements of MDA, SOD and CAT

GES-1 cells with good condition were cultured in 6-well plates. After the four groups described above are constructed, the cells were collected and counted, and detected by chemiluminescence according to the kit instructions. For *in vitro* cell experiments, the cells were collected into the centrifuge tube, and then the supernatant was discarded after centrifugation. 1 mL extraction solution was added for every 5 million cells and ultrasonic crushing was performed (power 200 w, ultrasonic 3 s, interval 10 s, repeat 30 times). Centrifuge 8000 g at 4 °C for 10 minutes, take supernatant and put it on ice for test. Gastric tissue of each rat was homogenized in freshly prepared ice cold 0.05□M potassium phosphate buffer (pH 7.4) with 20 times dilution. The supernatant was obtained by centrifuging the tissue homogenate at 15,000□×g at 4 °C for 10 min, which was used for the evaluation of the gastric MDA level, SOD and CAT activities, and the results were expressed as nmol/mg protein or U/mg protein. SOD activity was determined using a SOD assay Kit (BC0175, Solarbio, China); CAT activity was evaluated using a CAT assay (BC0205, Solarbio, China) and the MDA level was also determined by a MDA assay (BC0025, Solarbio, China).

### Mitochondrial staining and branch length analysis

After different treatments on GES-1 lived in Confocal dishes, PK Mito Orange (PKMO-1, Genvivo Biotech, China) and DAPI were added to the medium, followed by incubation at 37 °C for 15 min. The culture medium was then discarded and the cells washed gently three times with cell medium. Images were obtained under the Leica STELLARIS 5 Confocal Microscope Platform. Quantitative analysis of the mean branch length of mitochondria was calculated using MiNA software developed by ImageJ.

### Detection of mitochondrial membrane potential (MMP, △Ψm)

After the treatment, △Ψm of GES-1 cells were measured with a mitochondrial membrane potential detection kit (CA1310, Solarbio, China). In detail, after pretreatment with capsaicin for 2.5 h, cells were treated with 5% EtOH for 10 min or CCCP, a positive control reagent in the kit, for 20 min. Cells were collected and incubated in a cell incubator at 37 °C with JC-10 staining solution for 20 min. Then, we followed the instructions of the kit and analyzed it by flow cytometry.

### ATP detection

After pretreatment with capsaicin for 2.5 h, GES-1 cells were treated with 5% EtOH for 10 min, ATP in cell supernatant and intracellular were detected using an ATP test kit (S0026, Beyotime, China). The principle of the kit is that ATP is needed to generate fluorescence when firefly luciferase catalyzed luciferase. The fluorescence production is detected by the chemiluminescence instrument, which is proportional to the concentration of ATP within a certain range.

### Lactic acid (LA) detection

After CAP pretreatment for 2.5 h and EtOH incubation for 1.5 h, the intracellular LA level was measured using a Lactic Acid assay kit (A019-2-1, Nanjing Jiancheng Bioengineering Institute, China). Follow the kit instructions and test at 530 nm (EnSpire Multilabel Reader).

### Network targets analysis of CAP

Network targets analysis was performed to obtain the potential mechanism of effects of CAP on ROS^57–59^. Biological effect profile of CAP was predicted based our previous network-based algorithm:drugCIPHER^60–65^. Enrichment analysis was conducted based on R package clusterProfiler v4.9.1 and pathways or biological processes enriched with significant P value less than 0.05 (Benjamini-Hochberg adjustment) were remained for further studies. Then pathways or biological processes related to ROS and significantly enriched were filtered and classified into three modules, including ROS, inflammation and immune. Network targets of CAP against ROS were constructed based on above analyses.

### Proteomics and omics analysis

GES-1 cells were divided into three groups: Ctrl; EtOH and CAP + EtOH, and each group also had three biological replicates. After medium or 8 μM CAP pretreatment for 2.5 h and medium or 5% EtOH incubation for another 1.5h respectively, they were washed with PBS and 300 μl SDT lysis solution was added into a 10 cm petri dish. In order to obtain the differential expression genes (DEGs), this project adopted the quantitative proteomics technology of TMT, which mainly included two stages of mass spectrometry experiment and data analysis. The DEGs (Fold change ≥ 1.2, P value ≤ 0.05) were enriched by GO and KEGG through the DAVID database (https://david.ncifcrf.gov/), and the bioinformatic analysis was performed using the OmicStudio tools at https://www.omicstudio.cn/tool.

### Western blotting

At the specified time after processing, the cells and supernatant were collected directly into the centrifuge tube. After centrifugation at 6,500 rpm for 5 min, cells were washed three times with PBS and lysed in RIPA buffer. The protein samples were separated by 12% SDS-PAGE (80 V, 30 min and 130V, 55min), transferred to nitrocellulose(100 V, 105 min), sealed with 5% skim milk (30 min at RT), probed with the indicated primary antibodies (overnight at 4 °C), washed 3 times with TBST (120 rpm,6 min) and exposed to species-specific horseradish peroxidase (HRP)-conjugated secondary antibodies(60 rpm, 60 min). After washed 3 times with TBST, immunoreactive bands were visualized by enhanced chemiluminescence. The gray value of protein was analyzed by ImageJ software. GAPDH and ACTB served as the reference for equivalent total protein volume. WB of rat tissue was operated on ice at 4 °C during the whole process. The tissue was cut into fine fragments and the high efficiency RIPA tissue lysate (R0010, Solarbio, China) was added at the ratio of 200 µl lysate per 20 mg gastric tissue. The lysate was homogenized with a glass homogenizer until fully lysed. After centrifuging at 12000 rpm for 5 min, the supernatant was taken to conduct subsequent target protein detection.

### Quantitative PCR (RT-qPCR)

Total RNA from the cells was extracted using a QIAzol lysis reagent. The RNA was reverse-transcribed into cDNA using TransScript first-strand cDNA synthesis supermix according to the manufacturer’s instruction (AE301-02, TransGen Biotech, China). Hieff UNICON® Universal Blue qPCR SYBR Green Master Mix was used for qPCR (11184ES03, Yeasen, China). The relative amount of target gene mRNA was normalized to that of *GAPDH* mRNA in the same sample. The primers’ sequences used in cell experiments were designed through PrimerBank (https://pga.mgh.harvard.edu/primerbank/) and the primers’ sequences of PKM2 and LDHA were designed through a reference^66^. The sequence of qPCR primers in the rat experiments were listed using *r-ACTB* and etc. All the primer sequences of RT-qPCR were listing in **Table 1**.

**Table 1:**
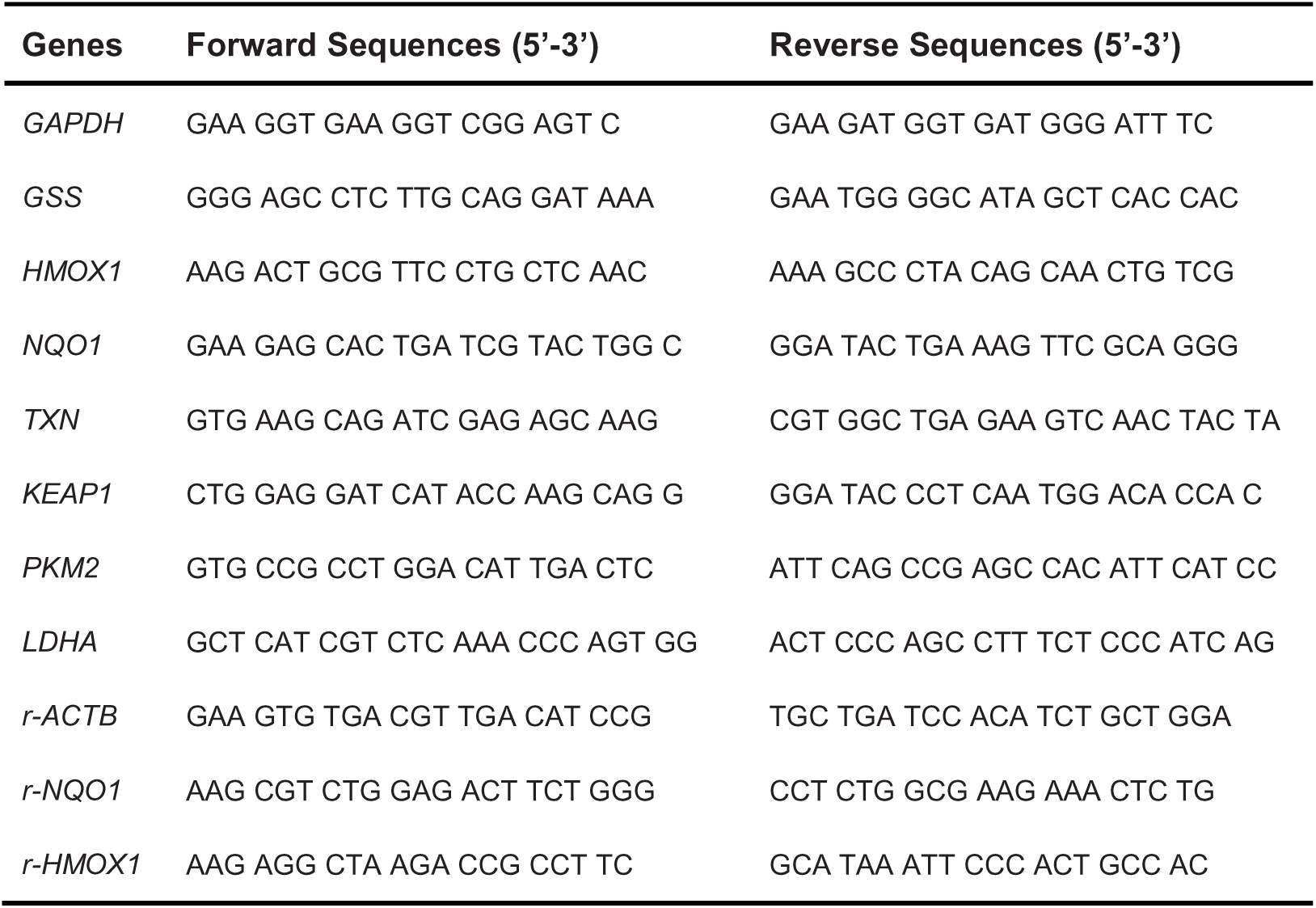
The primer sequences of RT-qPCR.

**Table 2:**
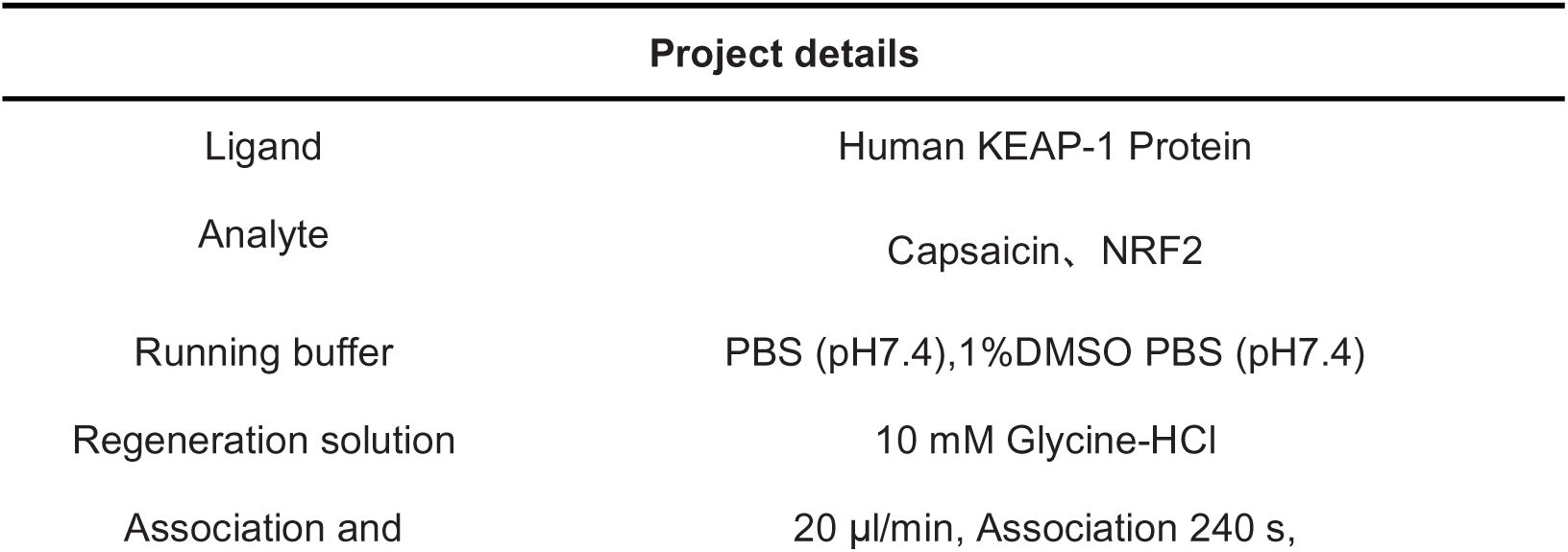
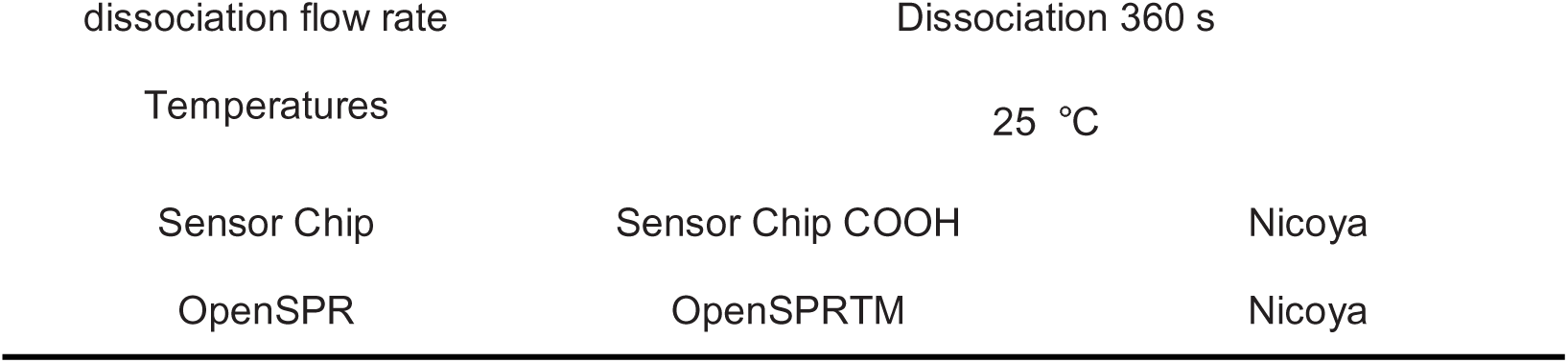
Experimental method design and parameters.

| Project details |  |
| --- | --- |
| Ligand | Human KEAP-1 Protein |
| Analyte | Capsaicin, NRF2 |
| Running buffer | PBS (pH7.4),1%DMSO PBS (pH7.4) |
| Regeneration solution | 10 mM Glycine-HCl |
| Association and | 20 μl/min, Association 240 s, |

|  |  |  |
| --- | --- | --- |
| dissociation flow rate | Dissociation 360 s |  |
| Temperatures | 25 °C |  |
| Sensor Chip | Sensor Chip COOH | Nicoya |
| OpenSPR | OpenSPRTM | Nicoya |

### Immunofluorescence staining and confocal microscopy

At the indicated times of treatment with CAP and EtOH, GES-1 cells were fixed in 4% paraformaldehyde (PFA) for 15 min, permeabilized with 0.5% Triton X-100 for 10 min, blocked with 1% bovine serum albumin (BSA) for 20 min, and then probed with specific antibodies (dilution ratio of the primary antibody of NRF2, 1:100; secondary antibody, 1:500). Then, 10 mg/mL DAPI was used to label nuclei. Images were captured with a confocal laser scanning microscope system. The colocalization of signals from two channels was detected using ImageJ software.

### Nucleoplasm separation

After different treatments on GES-1 cultured in 6-well plates, cells were centrifuged at 6000 rpm for 5 min and discarded the supernatant. All subsequent operations were performed on ice at 4 °C. First, the cells were decomposed for 5 min by adding 100ul of cytoplasmic lysate (containing 10 mM HEPES, 10 mM KCl, 0.1 mM EDTA, 0.5% NP-40 and adding extra 1:1000 PMSF and 1:2000 DTT). Then blow 200 times with a 200 µl Eppendorf. After centrifugation at 500 g for 5 min, the supernatant was the cytoplasmic lysate. Finally, 200 µl of NP40-free lysate was added for nuclear washing and precipitation for 3 times (discard the supernatant). After mixing with an oscillator, the mixture was centrifuged at 500 g for 5 min to obtain the nuclear proteins. All samples were boiled at 99 °C for 10 min for WB.

### Coimmunoprecipitation (Co-IP)

First, 3 mg pcDNA3.1 or Keap1-FLAG plasmid with polyethylenimine (PEI) reagent was transfected into 293T cells (5 × 10^6^ cells/ml). After 24 h, the cells were treated with 32 μM capsaicin for 2.5 hour and then suspended with phosphate-buffered saline (PBS) and centrifuged at 6,500 rpm for 5 min to collect cell precipitate. The cells were lysed in 700 ml lysis buffer (50 mM Tris-HCl, pH = 7.5, 150 mM NaCl, and 0.5% Triton-100), and the lysates were incubated with anti-FLAG affinity matrix at 4 °C for 10 h. The lysates were washed six times with PBS, eluted with glycine buffer (pH = 2), boiled in SDS loading buffer, and analyzed by WB. For the ubiquitin experiment, 3 mg Ub-K48-Myc and 3 mg VR1012 or NRF2-HA plasmids with PEI were co-transfected into 293T. After 36 h, the cells were treated with 32 μM capsaicin for 2.5 h. The rest of the steps were the same except that anti-HA affinity matrix was used.

### Pull-down

Purified Kelch-Strep protein (10 μg) was incubated with 10 μL of specific anti-Strep affinity matrix for a duration of 4 hours at 4°C. Solutions containing 32 μM and 100 μM CAP were prepared, and 1 μg of NRF2 protein was added to each reaction system. The reaction systems were further incubated at 4°C for another 2 hours. Subsequently, the unbound proteins were subjected to wash steps consistent with Co-IP procedures, followed by result assessment through Western Blot experiments.

### The degradation of NRF2

To test the degradation of NRF2, the drugs CHX and PS-341 were used. GES-1 cells were incubated with or without 8 μM CAP for 3 h, followed by treatment with CHX (50 μg/mL) for 15 min or 30 min. Total cell lysates were collected and analyzed by WB for total NRF2 protein expression. What’s more, GES-1 cells were incubated with or without PS-341(100 nM) and CAP (8 μM) for 3 h to compare the activation of NRF2 between co-incubation and single drug use. The total amount of NRF2 was quantified with ImageJ.

### Surface plasma resonance (SPR)

When the Human KEAP-1 Protein was fixed on the metal surface of the chip as a ligand protein, the mobile phase containing the analytical substance (NRF2 or Capsaicin) flowed through the stationary phase. The experiment was conducted in strict accordance with the relevant standard procedures of SPR. The analysis software used for the results of this experiment was TraceDrawer (Ridgeview Instruments ab, Sweden), and the analysis method was One to One analysis model. The design of experimental methods and parameters involved are shown in **Table 1**:

### Bio-layer interferometry assay (BLI)

The Octet RED96e system (Molecular Device, ForteBIO) is ideally suited for the characterization of protein-small molecule binding kinetics and binding affinity. The protein Kelch (WT) and Kelch (Mut) were purified, biotinylated and diluted to 50□μg/ml. The proteins were then immobilized on SSA sensors. The sensors were then blocked, washed and moved into wells containing various concentrations of the test compound of CAP in kinetic buffer. Program settings default to programs that bind proteins and small molecules in the system. The binding signals were identified and the results were analyzed using the OctetHT V10.0 software. R^2^ is an estimate of the goodness of fit with values close to 1 indicating an excellent fit.

### Cellular thermal shift assay (CETSA)

GES-1 cells on 24-well plates were treated with 1640 medium containing Capsaicin (8 μM) or 0.1%DMSO for 2.5-3 h. Cells were collected from each group and divided into seven PCR tubes on average. The cells were heated at the specified temperature (46-67 °C) for 3 min and cooled at room temperature. The cells were lysed with liquid nitrogen and freeze-thawed four times. The cell lysates were centrifuged at 12000 rpm for 30 min at 4 °C. The protein supernatant was analyzed by WB, and was quantified by ImageJ software.

### Mass spectrometry (MS-MS)

The final concentration of capsaicin was 50 μM (DMSO less than 0.5% by volume) after 20 μg Keap1 protein solution was added with 2 μl capsaicin (1 mM) and was dissolved in HEPES (pH=7.4). Incubate at 30 °C for 3 h in the dark. Separated by 10% SDS-PAGE gel (80 V 30 min and 130 V 20 min); The overexpressed KEAP1 was detected by Coomassie Bright Blue staining, and the corresponding bands were removed from the gel for intra-gel digestion as described. In short, the gel tablets were reduced with DTT (10 mM) and alkylated with iodoacetamide (50 mM). The rest of the process is followed and the digested protein samples are manipulated and analyzed by Novogene Co. Ltd., China.

### Computational simulations

The flexible docking between the KEAP1 Kelch domain and the small molecule CAP was performed by RosettaLigand^67^. The crystal structure of Kelch (PDB: 3WN7) and 3D model of CAP (ZINC1530575, https://zinc.docking.org/substances/ZINC000001530575/) were used as the input structures. The Rosetta software was obtained from https://www.rosettacommons.org and version number is 2021.36+release.57ac713 57ac713a6e1d8ce6f60269b3988b1adac1d96fc6. Rosetta Transform mover with the box size of 8.0 Å was used for global searching and the HighResDocker mover was used for fine tuning the binding positions. 1000 docking poses were generated and the interface binding energies was calculated using the default Ref2015_wts score functions. The docking results were ranked by the binding energy and clustered for the representative conformations.

The 100 ns all-atom simulations on the complex of NRF2-KEAP1 or NRF2-CAP-KEAP1 were performed with the Gromacs 2019.6 package^68^. The initial poses of inhibitor-binding conformations were adopted from the docking results and from PDB 3WN7. The system was solvated in a box (81 ‘ 81 ‘ 81 Å3) with TIP3P waters and 0.15 M NaCl. The CHARMM36 force field was adopted, wherein the topologies of inhibitors were generated by the CHARMM_GUI Ligand Modeler^69^. The energy minimizations were performed to relieve unfavorable contacts, followed by 10 ns equilibration steps. Subsequently, the simulations were performed at 300 K (velocity-rescale thermostat) and constant pressure (1 bar, Parrinello-Rahman NPT ensemble). The nonbonded interaction cut-off for electrostatics calculations was set as 10 Å and the particle mesh Ewald (PME) method was used for calculation of long-range electrostatic interactions. LINCS constraints were applied to h-bonds and the time step was 2 fs. Throughout the trajectories, the representative binding conformations were clustered based on their structural similarities.

### Protein purification

The synthetic cDNA encoding Kelch (WT) and Kelch (Mut) were cloned into pET28a vector respectively. The plasmid vectors were transformed into E. coli BL21 (DE3) to express proteins, which were induced by 0.4 mM isopropyl β-D-1-thiogalactopyranoside (IPTG) at 16°C for 18 hours, followed by cultured at 37 °C until the OD600 value reached 0.6. The cells were isolated by centrifugation, resuspended and lysed in lysis buffer. The lysates were centrifuged for 60 min at 12,000 rpm to obtain the supernatant. Gravity flow purification of the proteins in the supernatant were obtained using Ni-affinity chromatography and eluted in buffer containing 200 mM imidazole. The final proteins were purified and validated by Superdex 75 10/300GL resin.

### Hydrogen-deuterium exchange mass spectrometry (HDX-MS)

Kelch protein (1 mg/mL) was incubated with CAP (50 μM) or 0.2% DMSO in 4 °C overnight. The protein was equilibrated in D_2_O buffer in room temperature and quenched with ice-cold quench buffer (4 M Guanidine hydrochloride, 200 mM Citric acid and 500 mM TECP in water solution at pH 1.8, 100% H_2_O) at indicated times, then immediately put the sample tube on ice. Then, 1 μM pepsin solution was added for digestion. At 2 minutes, the sample was placed into a Thermo-Dionex Ultimate 3000 HPLC system autosampler for injection.

For LC-MS/MS analysis, the peptides were separated by a 20 min gradient elution at a flow rate 115 µl/min with a Thermo-Dionex Ultimate 3000 HPLC system, which was directly interfaced with a Thermo Scientific Q Exactive mass spectrometer. Peptides were separated on a reverse phase column (Acquity UPLC BEH C18 column 1.7 µm, 2.1*50 mm, Waters, UK). Mobile phase consisted of 1% formic acid, and mobile phase B consisted of 100% acetonitrile and 1% formic acid. The Q Exactive mass spectrometer was operated in the data-dependent acquisition mode using Xcalibur 2.0.0.0 software and there was a single full-scan mass spectrum in the orbitrap (350-2000 m/z, 70,000 resolution). The mass spectrometer was operated at a source temperature of 250 °C and a spray voltage of 3.0 kV. Peptic peptides were identified using an in-house Proteome Discoverer (Version PD1.4, Thermo-Fisher Scientific, USA).

The search criteria were as follows: no enzyme was required; two missed cleavage was allowed; precursor ion mass tolerances were set at 20 ppm for all MS acquired in an orbitrap mass analyzer; and the fragment ion mass tolerance was set at 0.02 Da for all MS2 spectra acquired. The peptide false discovery rate (FDR) was calculated using Percolator provided by PD. When the q value was smaller than 1%, the peptide spectrum match (PSM) was considered to be correct. FDR was determined based on PSMs when searched against the reverse, decoy database. Peptides only assigned to a given protein group were considered as unique. The false discovery rate (FDR) was also set to 0.01 for protein identifications. The deuterium exchange levels were determined by subtracting the centroid mass of undeuterated peptide from the centroid mass of deuterated peptide using HDExaminer (Version PD1.4, Thermo-Fisher Scientific, USA).

### Preparation of IR-HSA@CAP NPs

500 μl CAP (20 mg/ml) was uniformly injected into 10 ml of HSA solution (2 mg/mL), and stirred for 10 min at 700-800 rpm/min at RT. 50% glutaraldehyde was diluted 100 times and 80 μl was injected into the flask. Keep stirring at the same speed for 3 h. After termination of reaction, dialysis bag was used overnight to remove excess glutaraldehyde. HSA@CAP NPs were obtained and stored at 4 °C under shading. To obtain IR-HSA@CAP NPs, we added 40 μl of IRDye800 (5 mg/ml) into 700 μl of HSA@CAP NPs (2 mg/ml HSA). After incubation at RT for 1 h, the solution was added with a 30 kDa protein concentration tube, and centrifuged at 12000 rpm for 5 min. Some of the unbound antibody probes were centrifuged. Then eluted 4-5 times with 1XPBS (0.01 M) until there was no color in the collection tube. Protein concentration and labeling (DOL) were determined by measuring absorption at 280 nm and 780 nm.

### Particle size and Zeta potential

The morphology of the prepared IR-HSA@CAP NPs was observed by Transmission Electron Microscopy (TEM, TECNAI G2 F20). The sample was prepared as follows: IR-HSA@CAP NPs were diluted 150 times before ultrasonic treatment, so that the particles were evenly distributed in the solution. 10 μl sample solution was slowly dropped onto the surface of the copper mesh, covered with a protective cover to prevent dust pollution in the air, and waited for the sample solvent to fully volatilize at RT. The hydration particle size and potential of the samples were measured using Litesizer™ 500. 2-3 ml of aqueous solution diluted 1000 times by HSA@CAP and IR-HSA@CAP NPs were added into the plastic sample plate, and the hydration particle size and potential were detected at 25 °C. The same sample was measured at least three times in parallel to calculate the mean.

### High Performance Liquid Chromatography (HPLC)

The peak time and area corresponding to the HPLC were used to calculate the capsaicin content in IR-HSA@CAP. HPLC procedure: sample size was 10 μl, the temperature of C18 was adjusted to 30 °C, acetonitrile (10% - 94%) and 0.1% phosphoric acid water were used as mobile phase and time range was 0∼28 min. To evaluate the release of capsaicin in simulated gastric fluid (SGF:3.2 g pepsin, 2 g NaCl, 7 ml Concentrated hydrochloric acid in 1L ddH_2_0, pH=2.0), 200 μl of evenly mixed IR-HSA@CAP was added to 800 μl of freshly configured SGF/SPSS, and mixed at 37 °C. At the corresponding time point, it was evenly mixed with anhydrous methanol at 1:1 and added 1 ml mixed solution into an injection bottle for measurement.

### Endocytosis and *in vivo* imaging of IR-HSA@CAP NPs

*In vivo* experiments, healthy adult male Sprague-Dawley (SD) rats, weighing 230-260 g were used. The newly prepared IR-HSA@CAP NPs solution was diluted and added into the medium, and incubated with GES-1 cells for 1 h, 2 h and 3.5 h. After the supernatant was discarded, the cells were washed with PBS for 5 times, and the ICG fluorescence in the cells was detected by APC-Cv7-A channel of flow cytometry. SD rats (24 h after fasting) were given 1 ml IR-HSA@CAP NPs by intragastric administration. The fluorescence signals of rats were detected after administration. At the same time, images of side position were photographed at 45 min and 60 min, and dissected major organs (heart, liver, spleen, lung and kidney) and digestive system (stomach, large intestine, colon and duodenum) *in vitro* to determine the specific location of CAP.

### Pathological examination of gastric tissue (H&E Staining)

As previously mentioned, the stomach tissue is soaked in 4% paraformaldehyde tissue fixation solution for 24 h. Then the stomach tissue was cleaned, dehydrated by different concentrations of ethanol solution, transparent in xylene, and finally embedded in paraffin. Sections of 3 µm thickness were prepared, dried and dewaxed, stained with hematoxylin and eosin (H & E), dehydrated and sealed, and any structural changes were detected microscopically (OLYMPUS, DP26).

### Detection of cytokines in gastric tissue (ELISA)

Interleukin (IL)-1β, IL-6, CXCL1/KC(IL-8), IL-10 and tumor necrosis factor-alpha(TNF-α) levels in the gastric tissue lysate were determined to assess gastric inflammation. ELISA kits from Solarbio, catalogue numbers SEKR-0002, SEKR-0005, SEKR-0014, SEKR-0006 and SEKR-0009 were used for IL-1β, IL-6, CXCL1, IL-10 and TNF-α, respectively following the manufacturer’s instructions.

### Protocol for normal mice and Nfe2l2-KO mice

To ensure gastric emptying, mice were subjected to a 24-hour fasting period. Subsequently, they were administered SPSS or IR-HSA@CAP NPs. After a 45-minute pretreatment phase, the mice were exposed to 80% EtOH for an additional one hour. Following this, the animals were humanely euthanized, and the gastric tissue samples were collected for subsequent experiments and analyses. Precise dosages administered were as follows: IR-HSA@CAP nanoparticles used were prepared to contain 1 mg/kg of CAP and 2 mg/kg of HSA, as quantified by HPLC three times.

### Statistical analysis

The differences in the mean values between the two groups were analyzed using the student’s t-test, while the differences among multiple groups were analyzed using a one-way ANOVA test in the GraphPad Prism software. Therefore, we checked and proofread the statistical analysis. Statistical significant was indicated at a P value < 0.05. (ns, no significance; *, P < 0.05; **, P < 0.01; ***, P <0.001; ****, P <0.0001).

## Ethics statement

All animal research procedures were followed in accordance with the standards set forth in the eighth edition of Guide for the Care and Use of Laboratory Animals (published by the National Academy of Sciences, The National Academies Press, Washington, D.C.), and was approved by the Animal Care Committee of the Faculty of Medicine, Tianjin University (Approval No: TJUE-2023-238 and TJUE-2024-548).

## Acknowledgements

This work was financially supported by National Natural Science Foundation of China (22077094) and Key Research Project of the 14th Five-Year Plan for Cancer Prevention and Treatment at Tianjin Cancer Institute (No. YZ-08).

## Data availability

The data used to support the findings of this study are included within the article.

## Contributions

Xiaoning Gao: experiments, methodology, data analysis and writing-original draft; WuYan Guo: conceptualization and editing. Bo Zhang, Peiyuan Liu, Liren Liu and Zixiang Liu: conceptualization, methodology, data curation and resources; Mingyue Yuwen, Ruyang Tan and Junli Ba: experiments and visualization; Hongyu Ren and Shengtao Hu: experiments and writing-review and editing. Zhiru Yang, Xue Bai, Shama Shiti, Kai Miao, Haozhi Pei and Cong Tang: resources and data analysis; Kairui Liu: writing-polish; Tao Wang and Cheng Zhu: investigation, data curation and writing-review; Jun Kang: conceptualization; supervision; writing-original draft; funding acquisition; writing-review and editing.

