## Supplementary Figure for "Capsaicin acts as a novel NRF2 agonist to suppress ethanol induced gastric mucosa oxidative damage by directly disrupting the KEAP1-NRF2 interaction": Supplementary Figure.pptx

### Slide 1
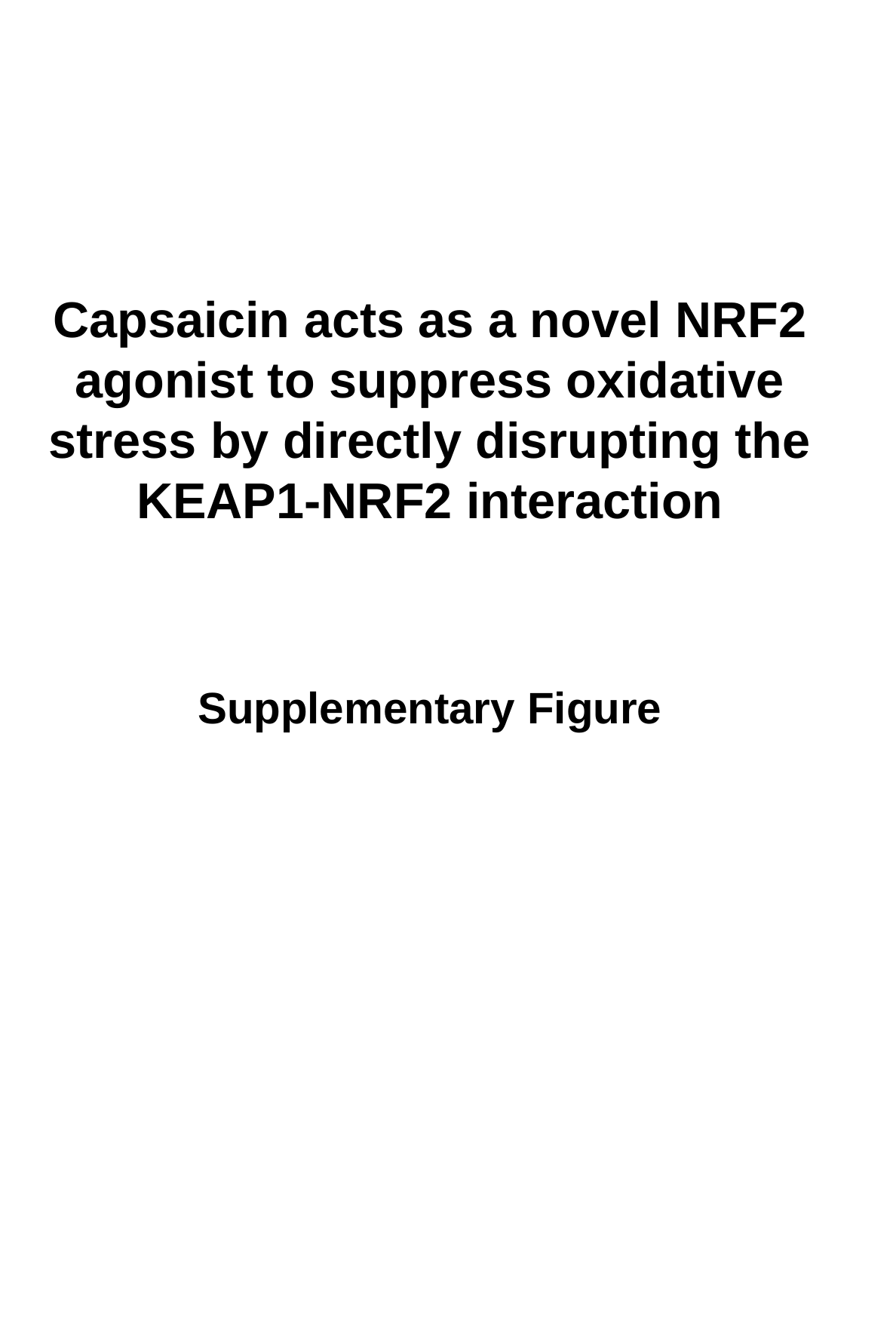

Capsaicin acts as a novel NRF2 agonist to suppress oxidative stress by directly disrupting the KEAP1-NRF2 interaction
Supplementary Figure

### Slide 2
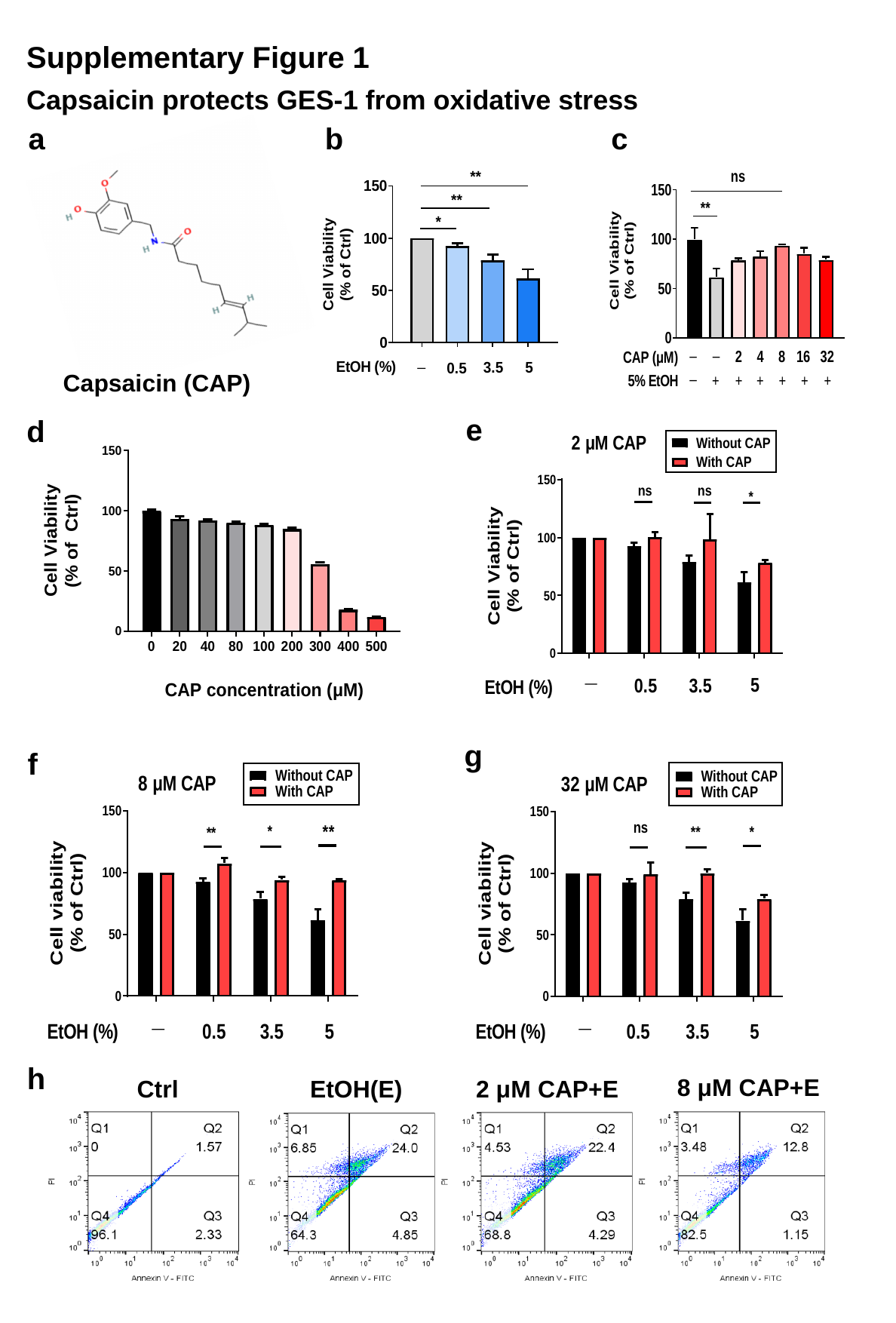

Supplementary Figure 1
Capsaicin protects GES-1 from oxidative stress
a
b
c
Capsaicin (CAP)
e
d
g
f
h
8 μM CAP+E
Ctrl
EtOH(E)
2 μM CAP+E

### Slide 3
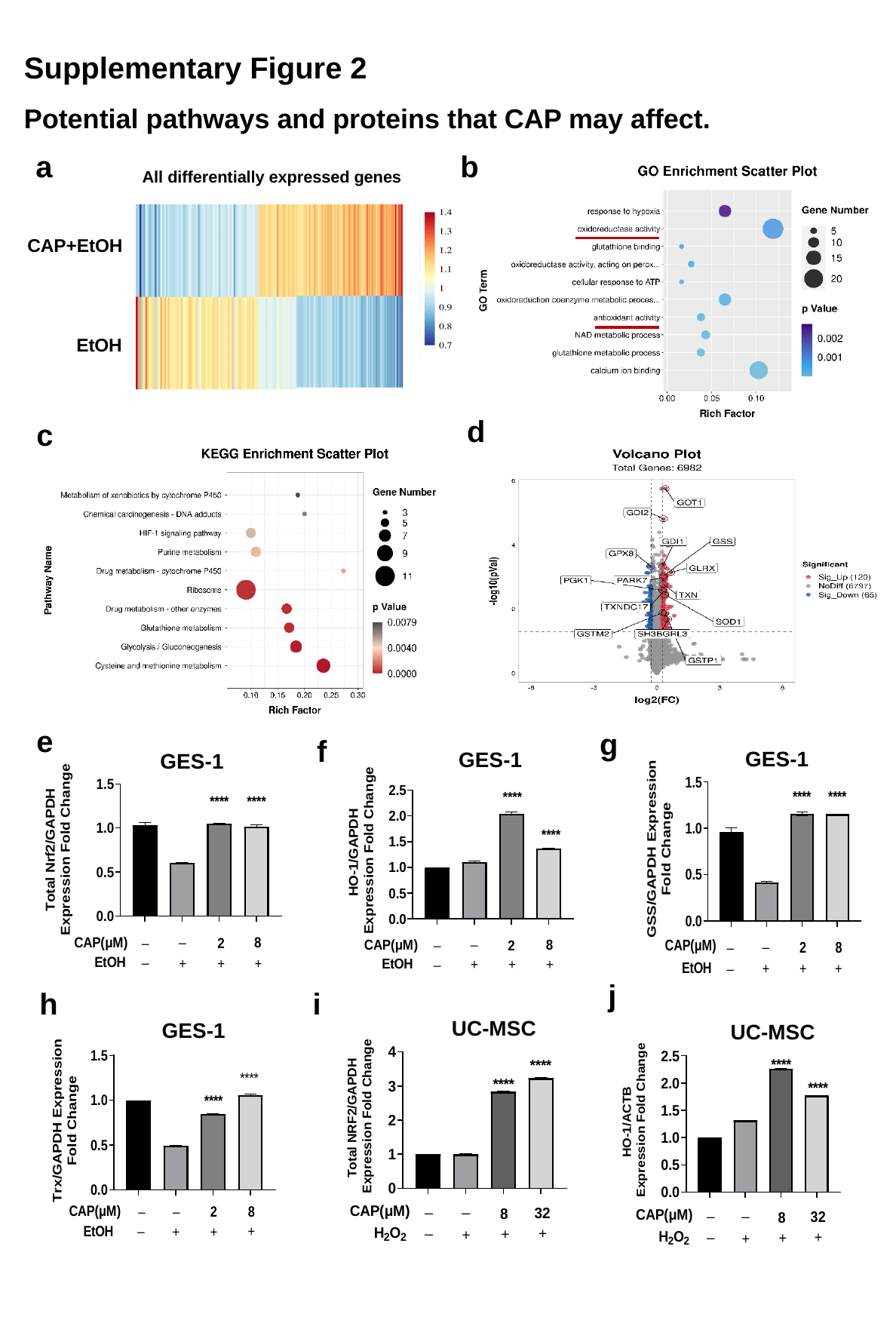

Supplementary Figure 2
Potential pathways and proteins that CAP may affect.
b
a
All differentially expressed genes
CAP+EtOH
EtOH
d
c
e
g
f
GES-1
GES-1
GES-1
j
h
i
UC-MSC
GES-1
UC-MSC

### Slide 4
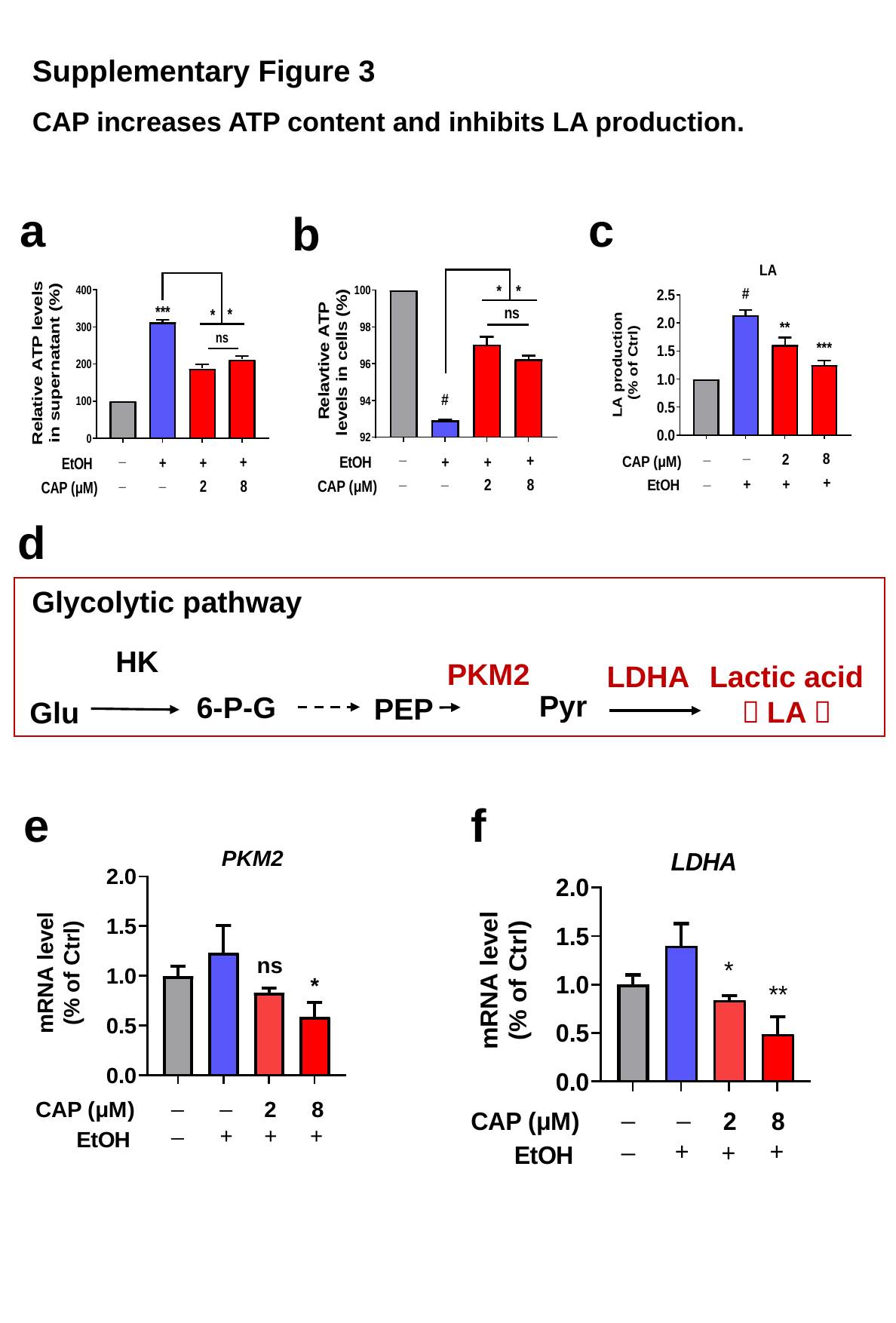

Supplementary Figure 3
CAP increases ATP content and inhibits LA production.
c
a
b
d
Glycolytic pathway
HK
PKM2
LDHA
Lactic acid （LA）
Pyr
6-P-G
PEP
Glu
f
e

### Slide 5
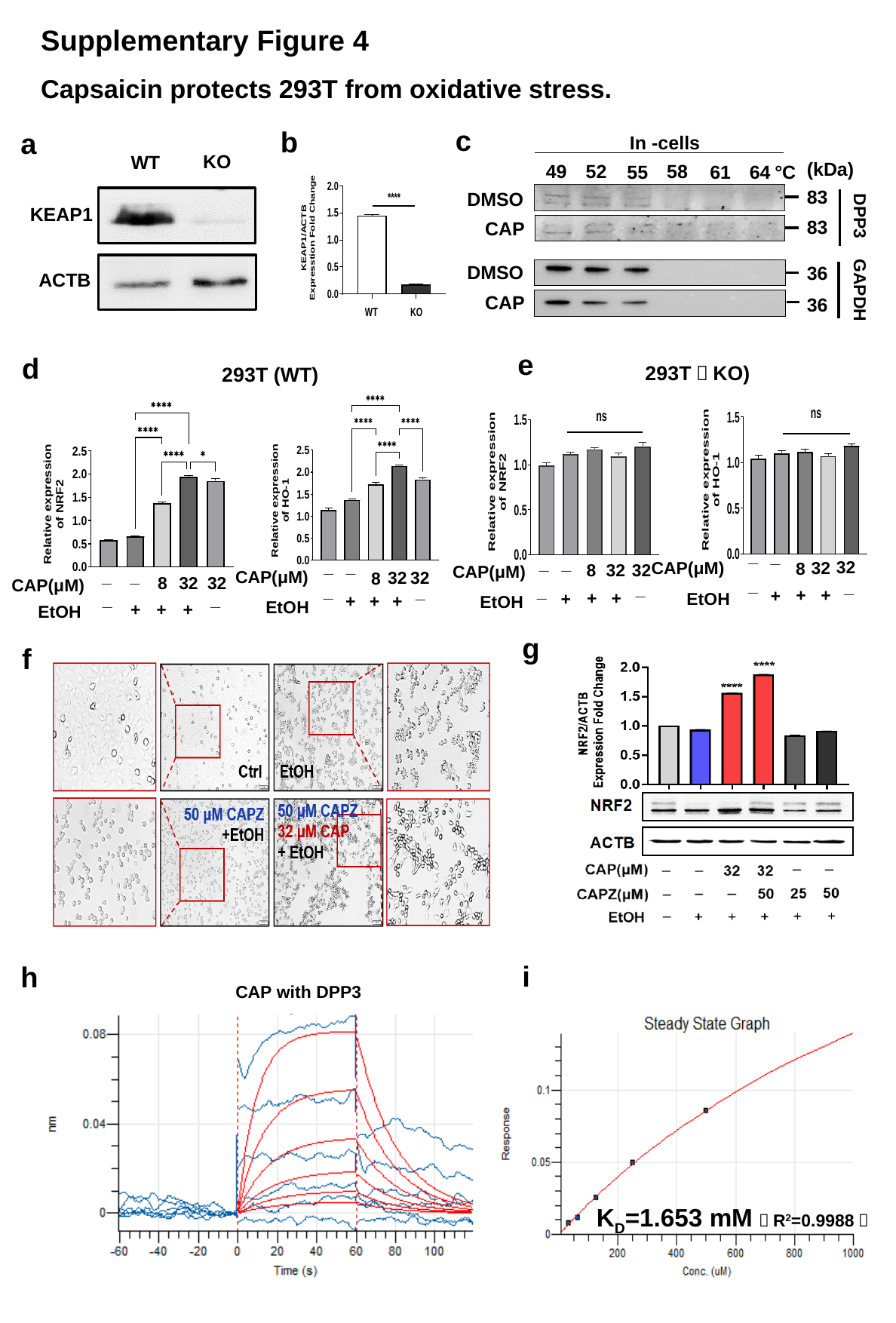

Supplementary Figure 4
Capsaicin protects 293T from oxidative stress.
c
b
a
In -cells
(kDa)
49
52
58
61
°C
55
64
83
DMSO
DPP3
83
CAP
DMSO
36
GAPDH
CAP
36
KO
WT
KEAP1
ACTB
e
d
293T（KO)
293T (WT)
_
_
CAP(μM)
32
32
8
_
_
+
+
+
EtOH
_
_
32
32
8
CAP(μM)
_
_
+
+
+
EtOH
_
_
32
32
CAP(μM)
8
_
_
+
+
+
EtOH
_
_
8
32
32
CAP(μM)
_
_
+
+
+
EtOH
g
f
i
h
CAP with DPP3
K_D_=1.653 mM（R^2^=0.9988）

### Slide 6
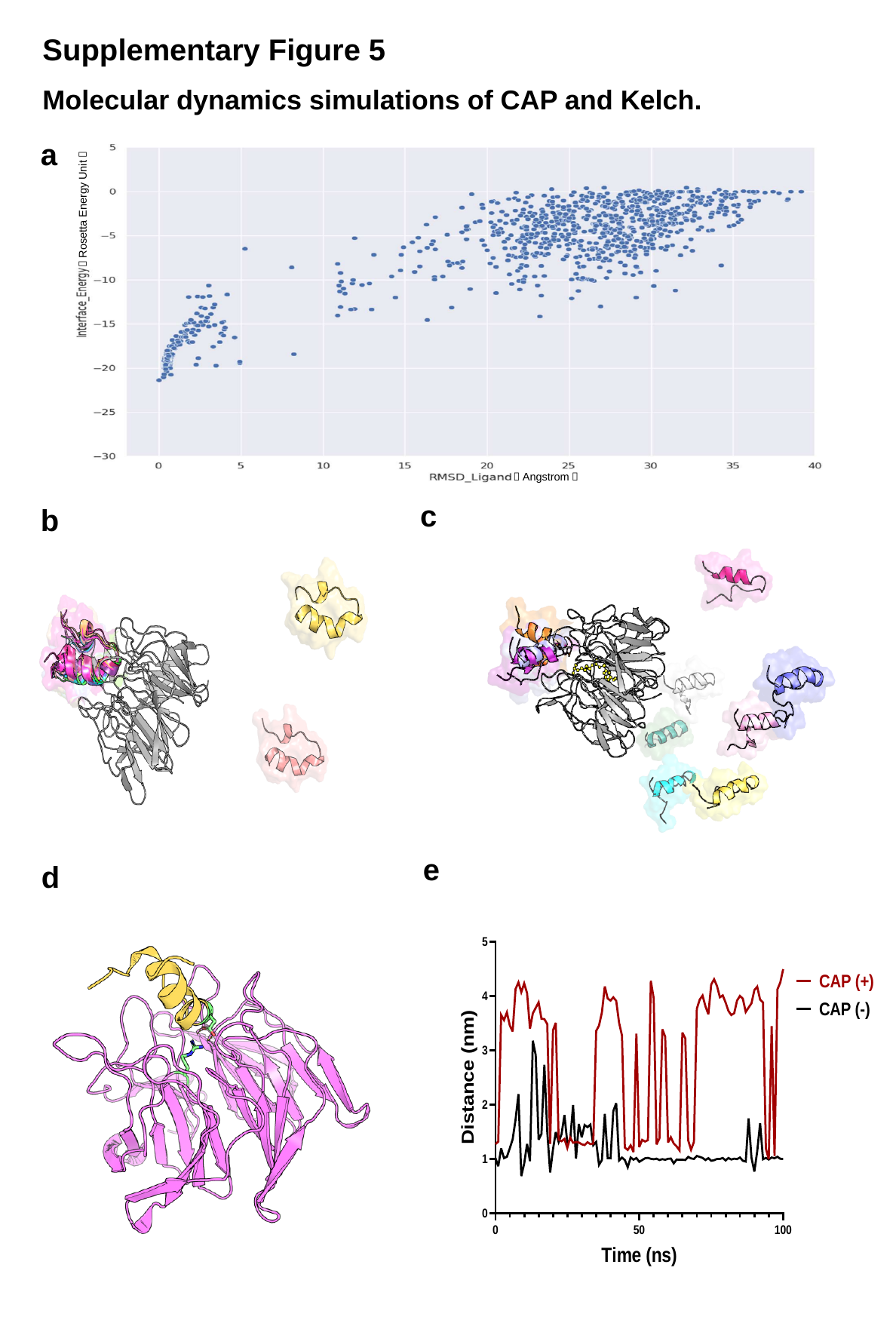

Supplementary Figure 5
Molecular dynamics simulations of CAP and Kelch.
a
（Rosetta Energy Unit）
（Angstrom）
c
b
e
d

### Slide 7
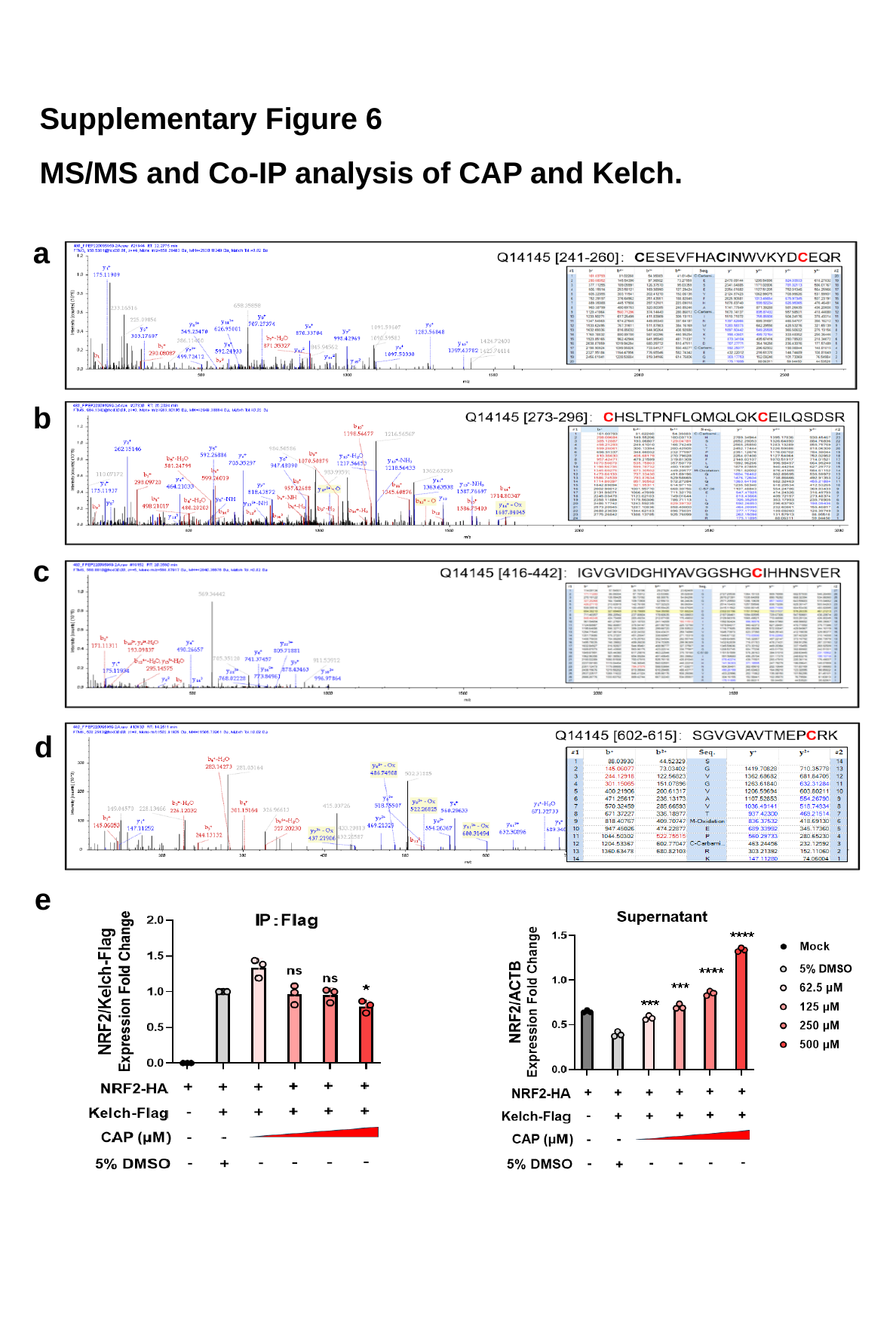

Supplementary Figure 6
MS/MS and Co-IP analysis of CAP and Kelch.
a
b
c
d
e

### Slide 8
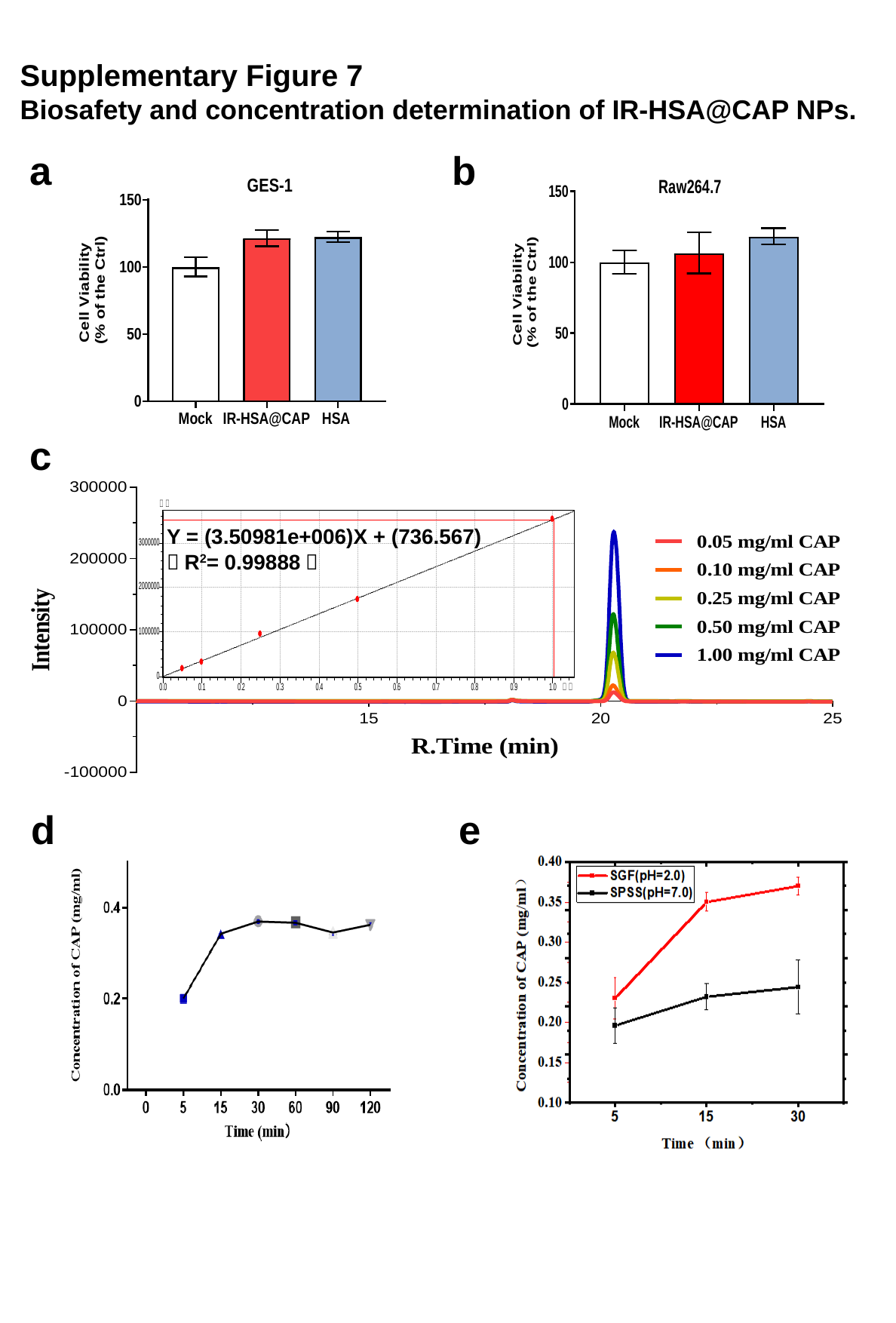

Supplementary Figure 7
Biosafety and concentration determination of IR-HSA@CAP NPs.
a
b
c
Y = (3.50981e+006)X + (736.567)
（R^2^= 0.99888）
d
e

### Slide 9
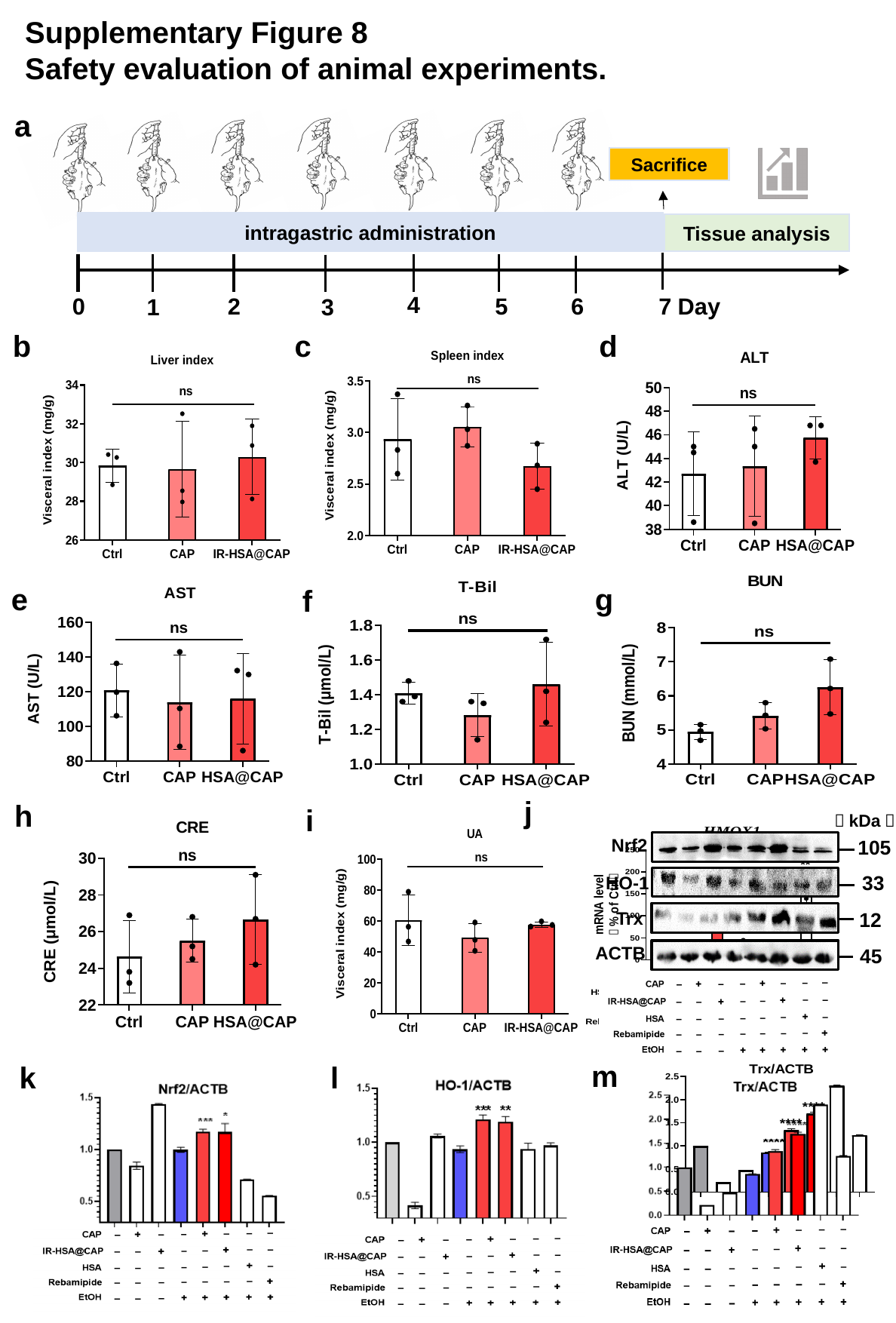

Supplementary Figure 8
Safety evaluation of animal experiments.
a
Sacrifice
intragastric administration
Tissue analysis
4
2
6
0
5
7 Day
3
1
b
c
d
g
e
f
j
h
i
（kDa）
Nrf2
105
33
HO-1
Trx
12
ACTB
45
m
k
l
