## Supplementary material for "Capsaicin acts as a novel NRF2 agonist to suppress ethanol induced gastric mucosa oxidative damage by directly disrupting the KEAP1-NRF2 interaction": Supplementary Materials.docx

**This PDF file includes:**

Figures. S1 to S7

Videos. S1 and S2 (MP4)

**SUPPLYMENTARY FIGURE LEGEND**

**Fig. S1 Capsaicin protects GES-1 from oxidative stress**

(a) Molecular formula of CAP from PubChem (Compound CID: 1548943). **(b)** Cell viability of GES-1 after 1.5 h treatment with ethanol at different concentrations (v/v). **(c)** Cell viability of GES-1 after pretreatment with different concentrations of CAP for 2.5 h and adding 5% ethanol for 1.5 h. **(d)** Effects of different concentrations of CAP on the viability of GES-1 cells for 24 h. **(e-g)** Cell viability was measured by adding CAP (2 μM, 8 μM or 32 μM) to GES-1 cells for 2.5 hours and then adding three concentrations of ethanol (0.5%, 3.5% and 5%) for another 1.5 hours. **(h)** FCM showed CAP inhibited the apoptosis of GES-1 induced by EtOH.

**Fig. S2 Potential pathways and proteins that CAP may affect.**

**(a)** Heat map showed the up-regulation and down-regulation of all differentially expressed proteins in CAP (8 μM,2.5 h) and 5% EtOH (1.5 h) versus 5%EtOH (1.5 h) alone. Red and blue are indicative of increased and decreased expression, respectively. **(b)** GO enrichment analysis of differentially expressed genes (DEGs). **(c)** KEGG enrichment analysis of DEGs. **(d)** Volcano plot illustrating DEGs between two treatment groups in GES-1 cells. Genes with significantly increased expression were marked in red, while those with significantly decreased expression were marked in blue. **(e-h)** Grayscale analysis of WB in GES-1. Statistical results of the expression of total Nrf2, HO-1, GSS and Trx compared with internal reference protein GAPDH. Experiments were repeated 3 times. **(i and j)** Grayscale analysis of western blot in HUC-MSC. Statistical results of the expression of total Nrf2, HO-1, GSS and Trx compared with internal reference protein GAPDH.

**Fig. S3 CAP increases ATP content and inhibits LA production.**

(a) The content of ATP in GES-1 cells in different treatment groups. **(b)** CAP (8 μM) pretreatment for 2.5 h and 5% EtOH incubation for 1.5 h, the intracellular LA level was measured. **(c)** Schematic diagram of intracellular glycolysis. **(d and e)** Detection of the mRNA level of *PKM2* and *LDHA* in different treatment groups.

**Fig. S4** **Capsaicin protects 293T from oxidative stress.**

**(a and b)** Expression of KEAP1 in human KEAP1 knockout 293T cells (293T KO). **(c)** Detection of DPP3-CAP interaction in GES-1 cells using cellular thermal shift assay coupled with western blotting (CETSA-WB). **(d)** The grey analysis of NRF2 and HO-1 of each group in 293T. **(e)** The grey analysis of NRF2 and HO-1 of each group in 293T(KO). **(f)** Microscopic examination of GES-1 cellular morphology following treatment regimens. Cells were pre-treated with Capsaicin (CAP) or Capsazepine (CAPZ) for 2.5 hours, followed by incubation with 5% EtOH for 1.5 hours. Scale bar, 10 μm. **(g)** Western blot analysis of NRF2 protein levels under varying concentrations of CAP and CAPZ, with or without EIOH treatment. **(h and i)** *In vitro* detection of DPP3 with CAP using BLI.

**Fig. S5** **Molecular dynamics simulations of CAP and Kelch.**

(a) The binding energy landscapes of the molecular docking of CAP and KEAP1. **(b)** NRF2 associated with Keap1 in the absence of CAP. Majority of frames throughout the 100 ns trajectory featured NRF2 binding with Keap1. **(c)** The distribution of NRF2 fragments around Keap1 (gray color) in the presence of CAP (represented as yellow ball/stick). The representative conformations of NRF2 throughout the simulations were superimposed on KEAP1. **(d)** The averaged distance between NRF2 Asp29 and Keap1 Arg415 (represented as green sticks) were measured and recorded in the MD simulations. **(e)** The distance between NRF2 Asp29 and KEAP1 Arg415 in the NRF2-KEAP1 complex (black line) and the NRF2-CAP-KEAP1 complex (red line).

**Fig. S6** **MS/MS and Co-IP analysis of CAP and Kelch.**

**(a)** MS/MS of KEAP1 peptide containing Cys257 after CAP treatment. **(b)** MS/MS of KEAP1 peptide containing Cys273 and Cys288 after CAP treatment. **(c)** MS/MS of KEAP1 peptide containing Cys434 after CAP treatment. **(d)** MS/MS of KEAP1 peptide containing Cys613 after CAP treatment. **(e)** The grey analysis of NRF2 in IP group and in the supernatant of each group.

**Fig. S7 Biosafety and concentration determination of IR-HSA@CAP.**

**(a and b)** IR-HSA@CAP NPs including HSA (0.04 mg/ml) and CAP (0.02 mg/ml) were added into GES-1 and Raw264.7 cells for 24 h to assess biosafety. **(c)** Standard curve for determination of CAP by HPLC. **(d)** The concentration of CAP released from IR-HSA@CAP NPs (including 0.4 mg/ml CAP) after a certain period of treatment in SGF by HPLC. **(e)** The concentration of CAP released from IR-HSP@CAP NPs (including 0.4 mg/ml CAP) was measured in SGF and SPSS for 30 min respectively.

**Fig. S8** **Safety evaluation of animal experiments.**

(a) A model diagram of safety trial about CAP and IR-HSA@CAP NPs in rats.

**(b and c)** Organ index of liver and spleen. ns, not significant. Data were presented as mean ± SD, P values were calculated using T-test. ns, not significant. **(d-f)** Determination of ALT, AST and T-Bil in serum to evaluate liver function. **(g-i)** Determination of BUN, CRE and UA in serum to evaluate renal function. Data were presented as mean ± SD, P values were calculated using T-test. ns, not significant. **(j-m)** Western blot analysis of antioxidant protein expression of NRF2, HO-1 and Trx.

**Supplementary Video S1:**

Molecular dynamics simulations between NRF2 and Kelch.

**Supplementary Video S2:**

Molecular dynamics simulations between NRF2 and Kelch with CAP.
